# Massively parallel base editing screens to map variant effects on anti-tumor hallmarks of primary human T cells

**DOI:** 10.1101/2023.12.13.571465

**Authors:** Zachary H. Walsh, Parin Shah, Neeharika Kothapalli, Gergo Nikolenyi, Shivem B. Shah, Giuseppe Leuzzi, Michael Mu, Patricia Ho, Sinan Abuzaid, Zack D. Brodtman, Neil Vasan, Mohammed AlQuraishi, Joshua D. Milner, Alberto Ciccia, Johannes C. Melms, Benjamin Izar

## Abstract

Base editing enables generation of single nucleotide variants, but large-scale screening in primary human T cells is limited due to low editing efficiency, among other challenges^1^. Here, we developed a high-throughput approach for high-efficiency and massively parallel adenine and cytosine base-editor screening in primary human T cells. We performed multiple large-scale screens editing 102 genes with central functions in T cells and full-length tiling mutagenesis of selected genes, and read out variant effects on hallmarks of T cell anti-tumor immunity, including activation, proliferation, and cytokine production. We discovered a broad landscape of gain- and loss-of-function mutations, including in *PIK3CD* and its regulatory subunit encoded by *PIK3R1, LCK*, *AKT1, CTLA-4* and *JAK1*. We identified variants that affected several (e.g., *PIK3CD* C416R) or only selected (e.g. *LCK* Y505C) hallmarks of T cell activity, and functionally validated several hits by probing downstream signaling nodes and testing their impact on T cell polyfunctionality and proliferation. Using primary human T cells in which we engineered a T cell receptor (TCR) specific to a commonly presented tumor testis antigen as a model for cellular immunotherapy, we demonstrate that base edits identified in our screens can tune specific or broad T cell functions and ultimately improve tumor elimination while exerting minimal off-target activity. In summary, we present the first large-scale base editing screen in primary human T cells and provide a framework for scalable and targeted base editing at high efficiency. Coupled with multi-modal phenotypic mapping, we accurately nominate variants that produce a desirable T cell state and leverage these synthetic proteins to improve models of cellular cancer immunotherapies.

## INTRODUCTION

Cellular cancer immunotherapies, such as adoptive T cell transfer (ACT) and Chimeric Antigen Receptor (CAR) T cell therapies, have become an important tool in the armamentarium for clinical cancer care in selected diseases(1). ACT of tumor-infiltrating lymphocytes (TILs) produces responses in a subset of patients with metastatic melanoma who have progressed on multiple prior lines of therapies, including immune checkpoint blockade (ICB). CAR-T cells are now widely used in the care of different leukemias and lymphomas, resulting in durable response, including in diseases that have traditionally been resistant to conventional therapies(2,3). However, most cancer patients do not benefit from cell therapies, and resistance may be due to cancer cell intrinsic changes (e.g., B2M loss, or CD58 loss/downregulation) or modulation of the cell products in the immunosuppressive tumor-microenvironment (TME)(3,4).

Recent studies indicate that intrinsic properties of T cells, which are used to produce such cell therapies, determine anti-tumor capacity and clinical success(5–7). TIL transfer, for example, is more successful when T cells have a progenitor-like cell state(5). Similarly, CAR-T products in which pre-infusion T cells expressed a less differentiated memory cell state were more likely to produce clinical responses in patients with large B cell lymphomas(6). Interestingly, single-point mutations in central signaling proteins (e.g., *mTORC*) or immune checkpoints (e.g., *CD28*)(8),(9,10) are sufficient to engender a potentially favorable cell state by modulating T cell receptor (TCR) signaling strength, memory phenotype and cell proliferation.

With the advent CRISPR-Cas9 technologies, there has been a series of efforts to genome-edit cell products with the goal to improve of T cell-based therapies, such as deletion of *PDCD1* or *CTLA-4,* or co-deletion of the endogenous *TCR* and *B2M* to reduce the likelihood for graft-versus-host-disease(11). Furthermore, critical studies have extended the powerful discovery tool of genome-scale CRISPR-Cas9 knockout screens to primary human T cells, allowing for unbiased identification of genetic drivers of T cell proliferation and survival(12). However, CRISPR-Cas9 mediated gene disruption has several biological and technical limitations, including unknown effects on allosteric protein-protein interactions and changes in the tumor-immune synapse(4,13), endogenous immunogenic potential of constitutively expressed foreign proteins (e.g. Cas9)(14), a high rate of aneuploidy events in perturbed cells(15) and limited tunability of endogenous and potentially favorable synthetic proteins.

Emerging base editing (BE) technologies can overcome several of these challenges and generate precisely engineered T cell products. CRISPR-dependent BE uses cytosine and adenine base editors (CBE and ABE) to induce site-specific deamination of cytosine and adenine, leading to C>T and A>G base transitions, respectively(16). Consequently, CBE and ABE can generate 64 distinct amino acid substitutions. Coupled with relaxed protospacer-adjacent motif recognition sites (NG instead of NGG) of single-guide RNAs (sgRNAs), BE enables mutagenesis of endogenous DNA across a wide range of the genome. However, broad application of these methods to primary human T cells has been limited to date due to low editing efficiency, scalability and cell toxicity ensuing from existing delivery methods.

Here, we overcome existing challenges through improved production, delivery, and scalability of BE and achieve unprecedented single and multiplex editing efficiency in primary human T cells, while finding optimal balances between genome editing and maintaining cell viability. We performed several large-scale BE screens in multiple donors under acute and chronic T cell activation conditions and found novel variants that improve T cell polyfunctionality across multiple donors. Through completely virus-free production and protein-free base editing of specific epitope-reactive human T cells that harbor base edits nominated from these screens, we engineered T cell products with superior, on-target tumor-lytic ability. Thus, this study paves the way for rapid, efficient, and safe application of base editing to improve existing and future cellular immunotherapies.

## RESULTS

### High-efficiency base editing using different delivery methods in primary human T cells

We first sought to establish and test methods for delivering adenine- and cytosine base editors (ABE and CBE, respectively) and single-guide RNAs (sgRNAs) to primary human T cells that result in high editing efficiency while maintaining cell viability, two major determinants in the successful development of cell-based therapies (**Fig. 1a**)(1). First, we devised a strategy using *in vitro* transcription (IVT) to generate high-quality, stable mRNA molecules with reduced immunogenicity encoding for the ABE or CBE **(Extended Data Fig. 1a,b)** and deliver these to T cells via optimized electroporation with an sgRNA targeting either a splice donor or acceptor site or a start codon mutation in *CD2* or *B2M,* respectively (**Fig. 1b,c**). The base editor mRNA is translated by the endogenous T cell ribosomal machinery to a fully functional synthetic protein. With this approach, we achieved high efficiency with all tested sgRNAs resulting in protein loss as measured by flow cytometry, ranging up to 99.5% with ABE (**Fig. 1d**). We also show that multiple base edits can be performed through simultaneous delivery of two sgRNAs targeting *CD2* and *B2M*, without loss of editing efficiency for either target (**Extended Data Fig. 1c**). Similarly, we achieved up to 85.9% editing efficiency using a CBE (**Fig. 1e**).

**Figure 1.**
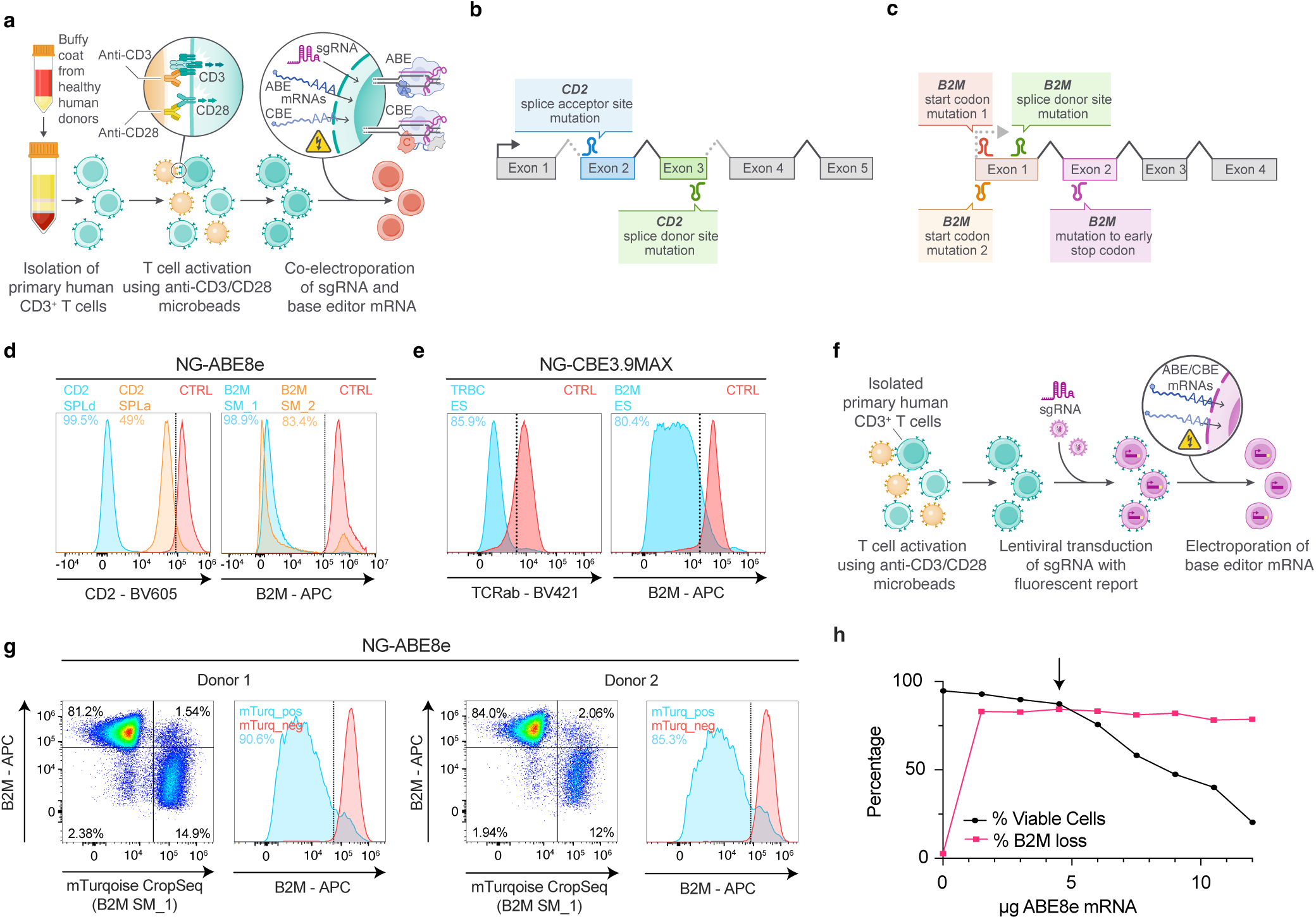
Establishment of methods for high-efficiency base editing in primary human T cells. **a,** Overview of approach for targeted base editing in primary human T cells. T cells isolated from deidentified human donors are CD3/CD28-stimulated for 48 hours, and then co-electroporated with a single guide RNA (sgRNA) and *in vitro* transcribed mRNA encoding the base editor (ABE = adenine base editor; CBE = cytosine base editor). ABE/CBE are translated to a functional protein in the target cell and performs sgRNA specified base edits. **b,c,** Target sites of sgRNAs against CD2 and beta-2-microglobulin (B2M) sites resulting in splice donor/acceptor mutations or mutations in start codons that results in early termination (SPLd = splice donor mutation, SPLa = splice acceptor mutation, SM = start codon mutation). **d,** Representative flow cytometry histograms showing ABE-mediated knockout of CD2 and B2M using sgRNAs indicated in (b) and (c). **e,** Representative flow cytometry histograms demonstrating CBE-mediated knockout of the T cell receptor (TCRab) and B2M (ES = early stop codon mutation). **f,** Approach for base editing primary human T cells with lentiviral integration of individual or pooled sgRNAs. T cells were isolated and activated as in (a), transduced with sgRNA (coupled with fluorescent mTurquoise reporter in a CropSeq lentiviral vector), and then electroporated with base editor mRNA 72 hours post-transduction. **g,** Representative flow cytometry dot-plots and histograms demonstrating ABE-mediated knockout of B2M with the workflow described in (f). mTurquoise fluorescence is used as a proxy for sgRNA integration. For histograms, red indicates gated mTurquoise-negative cells, and blue indicates gated mTurquoise-positive cells. **h,** Editing efficiency (measured by B2M loss on flow cytometry) and viability of T cells electroporated with B2M_SM_1 sgRNA and varying doses of the adenine base editor ABE8e. Timepoint is 4 days post-electroporation of ABE8e. Arrow represents ABE dosing selected for screening experiments.

To prepare for massively parallel BE screening, we tested another strategy: we delivered sgRNAs cloned into the CROP-seq (containing a mTurquoise fluorescent reporter) vector using lentiviral transduction followed by electroporation of BE mRNA **(Fig. 1f)**. We first validated the feasibility of this approach with CRISPR-Cas9 knockout, with lentiviral delivery of a previously validated sgRNA targeting *CD2* followed by nucleofection of SpCas9 mRNA **(Extended Data Fig. 1d)**. We then extended this approach to both ABE and CBE delivery and found unprecedented high efficiency in introducing start site mutations in *B2M* in two independent donors using ABE (90.6% and 85.3%) (**Fig. 1g**). The editing efficiency was essentially equal in both CD4+ and CD8+ T cells (**Extended Data Fig. 1e**). Furthermore, using a CBE we achieved high editing efficiency (56.5%) of the *TRBC* locus (**Extended Data Fig. 1f**). Importantly, we determined dosing of BE mRNA delivered to produce an optimal balance between high editing efficiency and cell viability (**Fig. 1h**), which is a prerequisite for performing large-scale screens in primary human T cells. In summary, we established a scalable strategy for high-efficiency base editing which enables base editor screening in primary human T cells.

### Massively parallel base editing to map variants to hallmarks of T cell function

To characterize variants broadly and deeply across a wide range of genes implicated in T cell function we designed two ABE sgRNA libraries. In the first library (“ClinVar library”), which included 8,142 sgRNAs, we mutagenized 102 genes involved in all major functions of T cells (**Fig. 2a, Extended Data Fig. 2a, Extended Data Table 1**), including known variants associated with clinical immune syndromes, variants of unknown significance (VUS), and other uncharacterized mutations that we generated at these defined loci. In the second library, we systematically tiled 7,815 sgRNAs to introduce every possible ABE-mediated mutation across the entire coding sequence and exon-intron boundaries of twelve genes with central functions in T cell activity, including T cell receptor signaling (*ZAP70, CD3Z, LCK, LAT*), costimulatory signaling (*CD2, CD28*), cytokine receptor signaling (*IL2RG, IL7RA, JAK1, STAT3, STAT5B*), and stemness (*TCF7*) (**Fig. 2b, Extended Data Fig. 2b**). Each library was designed to include multiple subtypes of control sgRNAs: Negative controls included empty-window sgRNAs (i.e. sgRNAs in which the defined base-editing window does not contain a targetable base), sgRNAs predicted to produce only silent mutations in our target gene set, and sgRNAs tiling a gene not involved in T cell function (*PPP1R12C*). Positive control sgRNAs were designed to generate an array of presumed-deleterious mutations, such as splice site mutations and missense mutations resulting in substitutions to proline, across 50 essential genes encoding factors critical for cell survival, including DNA polymerases (e.g. *POLR3C, POLR2E),* RNA splicing (e.g. *SF3B3)*, and cell-division (e.g. *KIF11*) **(Extended Data Fig. 2a,b, Extended Data Table 2)**. In all experiments, we used a highly active adenine base editor (ABE8e) with a relaxed protospacer-adjacent motif requirement (NG rather than NGG)(17), thus, maximizing the number of mutations that can be introduced.

**Figure 2.**
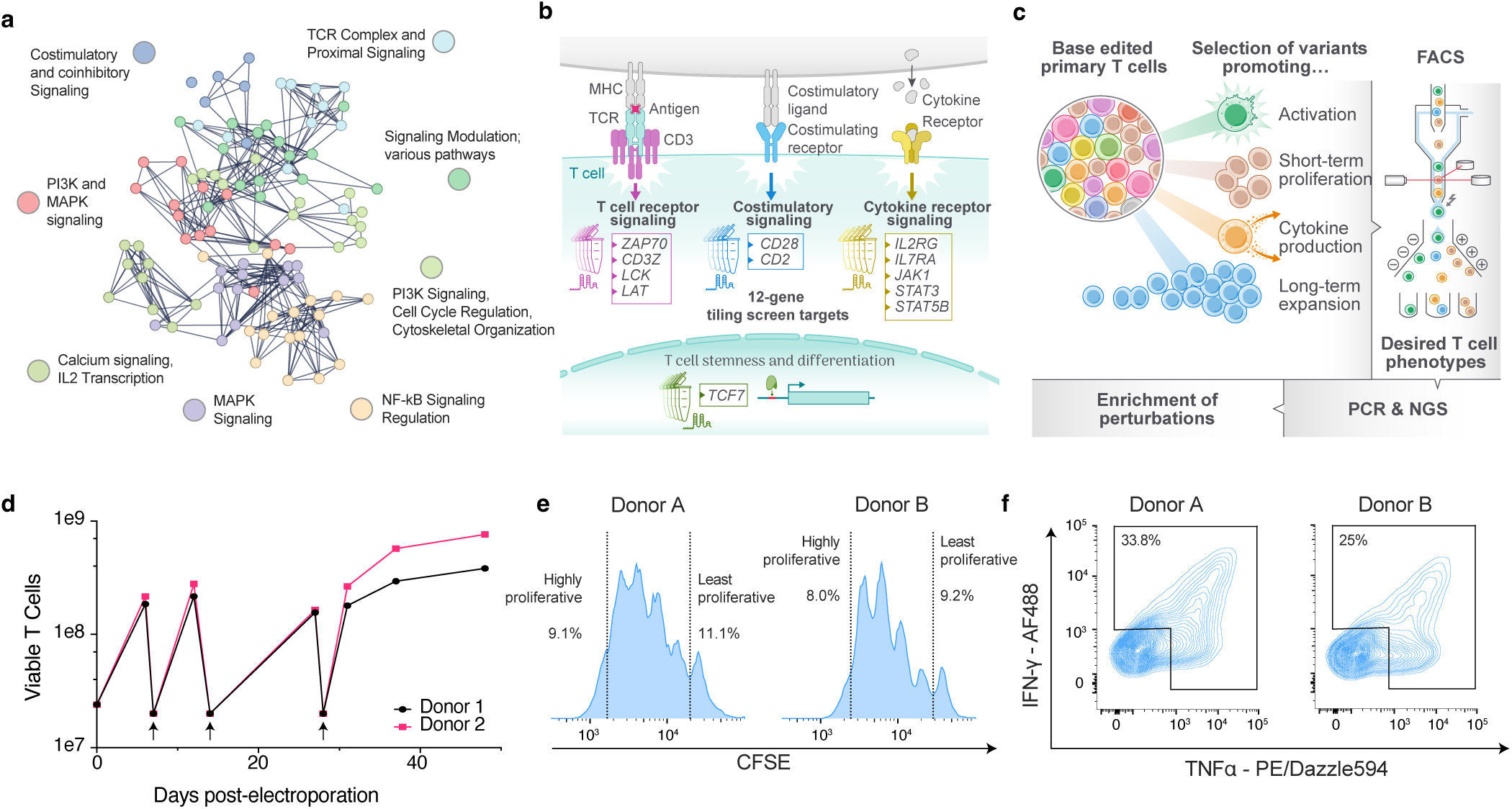
Design and execution of massively parallel base editor screens in T cells. **a,** Target gene set for ClinVar library ABE screens. Genes were selected from the “T cell receptor signaling” KEGG pathway, excluding secreted proteins. Visualization was performed in STRING with *k*-means clustering, *n*=8 clusters. Lines represent functional interactions. All sgRNAs generating either known ClinVar variants, or different point mutations at the same amino acid residue, were selected for the final library. **b,** Target gene set for 12-gene tiling library ABE screens. Genes were selected on the basis of their central functions in T cell signaling and differentiation. The sgRNA library was designed to tile the exons and splice sites of each gene to generate every possible ABE-mediated edit. **c,** Library-edited primary T cells were isolated (using FACS or serial collections of repeatedly stimulated T cells) for desired phenotypes and functions. Enrichment and depletion of guides in these readouts was assessed by next-generation sequencing (NGS). **d,** T cell numbers following base editing and chronic restimulation and long-term expansion arm of the ClinVar library screen. Arrows represent T cell restimulation points. At each restimulation point, 20e^6^ T cells were carried forward and restimulated with CD3/CD28 beads. T cells were sampled for NGS at days 7, 12, 15, 28, 35, and 40 following initial base editor electroporation. **e,** Representative histograms from the carboxyfluorescein succinimidyl ester (CFSE) short-term proliferation screen arm. Cells were sorted from the dimmest (most proliferative) and brightest (least proliferative) CFSE populations and sgRNA distribution was assessed by NGS. **f,** Representative contour plots from the cytokine production screen arm. T cells were CD3/CD28 stimulated for 6 hours and sorted for presence (either single- or double-positive) or absence (double-negative) of TNFα and IFNγ. Sorted populations were compared by WGS.

Following lentiviral transductions of these sgRNAs (cloned into the CROP-seq vector using optimized Golden Gate assembly, Methods) (**Extended Data Fig. 2c-d**) into pre-stimulated primary human CD3+ T cells isolated from two healthy donors, we electroporated ABE mRNA and allowed cells to edit and then proliferate for at least 7 days **(Extended Data Fig. 2e)**. The T cells from each screen were then probed with various assays to capture multiple critical axes of T cell function **(Fig. 2c)**, including activation, short- and long-term proliferation, and cytokine production. To approximate acute and chronic T cell receptor (TCR) engagement as seen in infections or cancer, respectively, base edited T cells were stimulated either through a single-time activation or with repetitive stimulations using CD3/CD28 microbeads (**Fig. 2d**). To assess edits affecting short-term proliferation capacity, cells were stained with Carboxyfluorescein Diacetate Succinimidyl Ester (CFSE), which enables tracking of cell proliferation by dye dilution, bead-stimulated, and allowed to proliferate for 4 days followed by flow sorting (CFSE^low^ vs. CFSE^high^) **(Fig. 2e)**. To determine variant effects on cytokine production, cells were briefly stimulated and sorted on intracellular presence of the proinflammatory cytokines IFNγ and TNFα (**Fig. 2f**). Finally, to assess edits impacting T cell activation status, we sorted T cells with high or low expression of the IL2 receptor alpha chain (CD25) **(Extended Data Fig. 2f)**. The distribution of sgRNAs in each condition was read out using next-generation sequencing (Methods).

### Base editor screens recover known variants critical for T cell survival and proliferation

We first analyzed several built-in controls across all experiments. In the ClinVar screen, we determined the distribution of sgRNAs targeting our gene set with either empty editing windows or generating silent mutations only, which showed no significant change in distribution at late timepoints in the long-term proliferation arm (**Fig. 3a, Extended Data Fig. 3a,b**), thus suggesting relatively low off-target base edit events. In contrast, sgRNAs disrupting essential genes (**Extended Data Table 2)** through either splice donor or splice acceptor splice site edits or, base edits resulting in amino acid changes to a proline (thus altering the structural organization of resulting gene products) were strongly depleted (**Fig. 3a,b, Extended Data Fig. 3a**). We noted that across both donors, sgRNAs generating splice donor disruption in essential genes had greater dropout than those generating splice acceptor disruption, consistent with previous reports indicating that splice donor mutations more reliably impact gene function(18). sgRNAs generating deleterious mutations in CD3 complex genes (*CD3D, CD3E*, *CD3G,* and *CD3Z),* which are essential for T cell activation, were significantly depleted across both donors **(Fig. 3a,b, Extended Data Fig. 3a)**. Finally, when assessing gene-wise dropout in the CD25^hi^ (activated) and CFSE^lo^ (highly proliferative) flow-sorted populations of the ClinVar screen, we observed depletion of variants in expected targets, including multiple members of the CD3 complex and genes essential for cell division **(Extended Data Fig. 3c,d)**.

**Figure 3.**
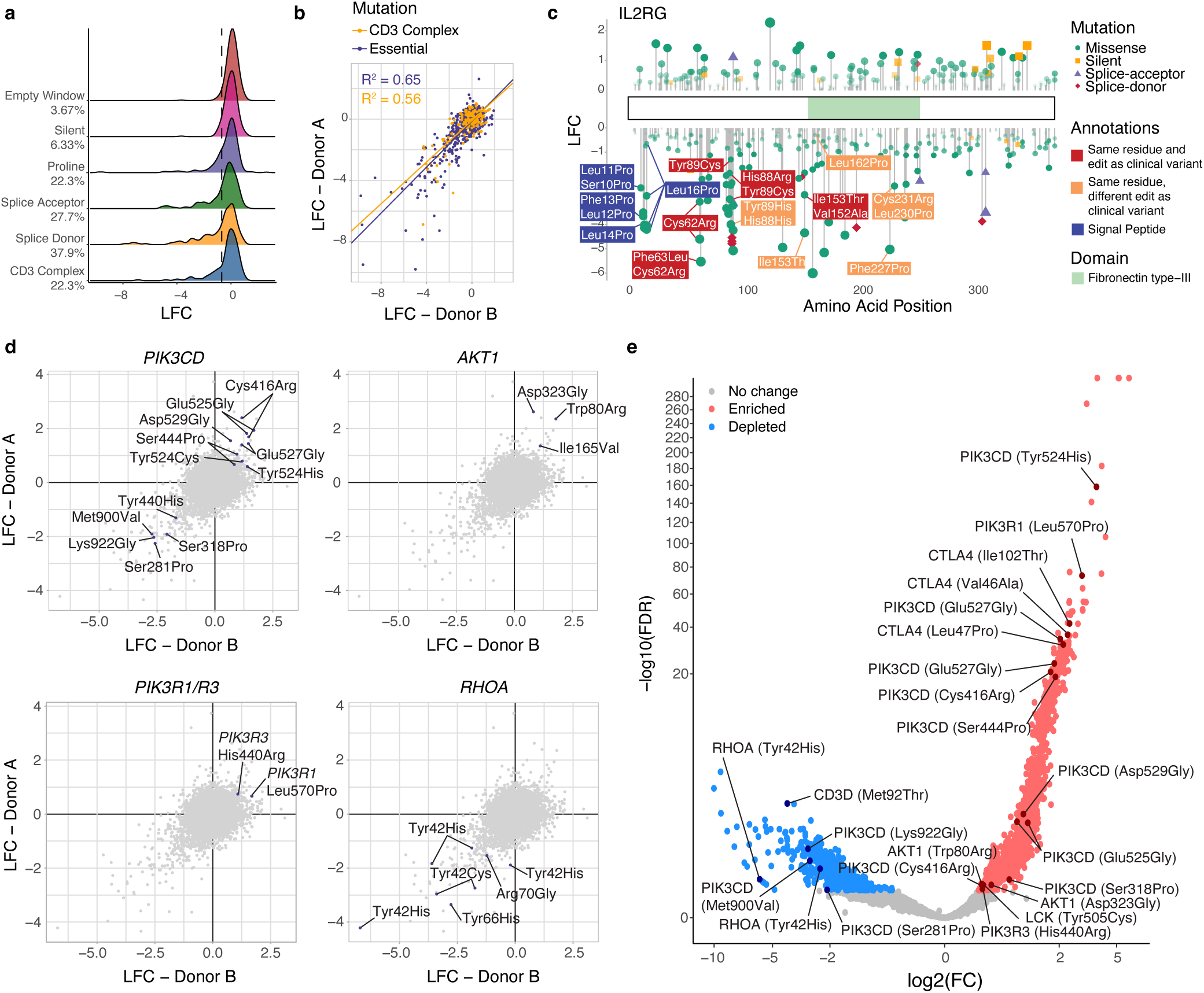
Global results of base editing screens across libraries, donors, and readouts. **a,** Density plots showing log2 fold changes (LFC) values of different categories of guides from the ClinVar library at Day 35 post-electroporation of the long-term expansion screen arm. Dotted line represents the bottom 5% of combined empty window and silent mutation controls. Percentages represent the percentage of guides in each category falling below this threshold. All guides generating variants in *CD3D, CD3E, CD3G,* or *CD3Z* were binned into the “CD3 complex” category. One representative donor is shown. SgRNA distributions from all long-term proliferation timepoints are compared to a “Day 0” control T cell population which was library-transduced but not electroporated with base editor. **b,** Scatter plot showing LFC values of essential gene or CD3 complex gene perturbations in both donors from the ClinVar Library at Day 28 post-electroporation in the long-term expansion screen arm, indicating high degree of robustness of the screens across two independent donor input T cells. **c,** Lollipop plot showing LFC of sgRNAs tiling *IL2RG* in the 12-gene tiling screen, mapped to the canonical IL2RG isoform. Indicated types of enriched and depleted mutations introduced by sgRNAs and their position in the amino acid sequence are shown. Selected sgRNAs generating clinically identified immunodeficiency variants (red boxes), alternative mutations at amino acids with known clinical variants (orange boxes), or variants in the signal peptide (blue boxes) that likely interfere with shuttling of IL2RG protein to the cell membrane are annotated. Timepoint shown is Day 26 post-nucleofection in the long-term expansion screen arm. **d,** Scatter plot showing LFC of sgRNAs across both donors (x vs y axis) in the long-term expansion ClinVar screen arm. Timepoint shown for all panels is Day 28 post-electroporation. Highlighted are positions of sgRNAs resulting in indicated amino acid changes for selected target genes and protein products (PIK3CD, AKT1, PIK3R1/3 and RHOA). **e,** Volcano plot showing LFC (x axis) and –log_10_(FDR) (y axis) for sgRNAs in the CFSE short-term proliferation arm of the ClinVar screen. Positive LFC represents enrichment (red) or depletion (blue) of the sgRNA in the highly proliferative population compared to the least proliferative. Indicated are selected amino acid changes in gene products from highly enriched/depleted base edits. FDR cutoff <0.05. Data from one representative donor is shown.

*IL2RG* encodes for the gamma chain of the IL2 receptor complex, whose ligand interleukin-2 (IL2) is required for T cell proliferation. In the 12-gene tiling screen, sgRNAs generating mutations at *IL2RG* splice-sites or introducing proline conversions (e.g. L16P, L14P, S10P) in the signal peptide, which is responsible for plasma membrane trafficking, were strongly depleted following multiple rounds of T cell restimulation (**Fig. 3c, Extended Data Fig. 3e)**. We also observed strong depletion of known *IL2RG* mutations (e.g. C62R, Y89C, L230P) that have been documented in patients with X-linked severe combined immunodeficiency (X-SCID), suggesting that base editor screens can recover and characterize variants of clinical significance(19). These results were highly consistent between both donors, indicating reproducibility of these screens **(Fig. 3c, Extended Data Fig. 3e)**.

Conversely, we also identified rational enrichment of sgRNAs in our screens reflecting mutations with known clinical, biological, and structural significance. In the long-term proliferation arm of the ClinVar screen, we observed enrichment of an sgRNA generating a mutation in the inhibitory histone domain interface of SOS1 (W85R), a guanine exchange factor which positively regulates Ras pathway signaling **(Extended Fig. 3f)**. W85R is a pathogenic mutation associated with Noonan Syndrome(20), a RASopathy driven by hyperactive Ras pathway signaling. Preclinical structural and functional studies have demonstrated that mutating this residue facilitates a partial release of SOS1 auto-inhibition, thus, increasing SOS1 signaling. Additionally, we found enrichment of a sgRNA predicted to generate a D323G mutation in AKT1, a likely-pathogenic mutation associated with familial breast cancer (**Fig. 3d**). Clinical and preclinical studies have shown that another missense mutation in the same residue – D323H – is a gain-of-function mutation driving increased downstream signaling, resistance to pharmacological allosteric inhibition(21). Moreover, the same preclinical studies identified a similar AKT1 variant (W80R) of unknown significance, which enriched in our long-term proliferation screen (see below) (**Fig. 3d**).

Together, these data suggest that we achieved excellent on-target base editing recovering enrichment and depletion of several expected gain- and loss-of-function mutations, respectively, while synonymous or empty-window base edits showed no differential abundance, suggesting low off-target base edits.

### Variants modulating T cell proliferation, activation, and effector function

Having established the robustness and reproducibility of the screens, we next quantified sgRNA abundance across multiple functional read-outs and identified variants modulating T cell proliferation, activation, and effector function (**Figure 3d-f, Extended Data Fig. 3f-l, Extended Data Fig 4-5, Extended Data Table 3-4**). In the long-term proliferation assay, we found significant enrichment of sgRNAs introducing an array of base edits in genes encoding protein kinases (*PIK3CD, PIK3R1, PIK3R3, AKT1, LCK),* phosphatases *(PTPRC,* which encodes CD45), and immune checkpoints (*CTLA4*), among others, as well as depletion of perturbations in genes encoding GTP binding proteins (*RHOA)* (**Fig. 3d, Extended Data Fig. 3f-j)**. We discovered clusters of highly enriched *PIK3CD* variants in the C2 domain (C416R, S444P) and helical domain (Y524H, Y524C, E525G, E527G, D529G) **(Fig. 3d)**. Other highly enriched variants involved the pleckstrin homology and protein kinase domains (W80R and I165V, respectively) of AKT1, the C-terminal tyrosine residue of LCK (Y505C), and the tyrosine phosphatase 1 domain of CD45 (N760S) **(Fig. 3d, Extended Data Fig. 3f)**. SgRNAs generating variants of RHOA in the effector domain (Y42H, Y42C) and switch II domain(22) (Y66H, R70G) were among the most significantly depleted across multiple donors **(Fig. 3d, Extended Data Fig. 3g,h)**. For many of these variants, we observed concordant enrichment/depletion patterns in the CFSE-proliferation assay (**Fig. 3e, Extended Data Fig. 3i)**. This suggests that these variants, particularly involving *PIK3CD*, have the potential not only for short-term, but sustained proliferative capacity, which is a prerequisite for robust T cell antitumor activity. We next identified putative variants associated with increased cytokine production, including in several genes with central functions in TCR signaling, co-stimulation, signal transduction and transcription regulation, such as ITK (Y588C, Y578H), ZAP70 (I203V), NFKB1 (E439G, N846D, E63G), CARD11 (I1076T), CTLA4 (D153G), and CD40LG (Q220R, Y172H, among others (**Extended Data Table 3**). In the same readout, we identified dozens of known or likely pathogenic variants in several genes resulting in a range of immunodeficiency disorders (**Extended Data Table 3**). Additionally, variants in genes identified in other readouts (e.g. *PIK3CD* and *CTLA4*) were also enriched in the cytokine assay (**Extended Data Table 3**). Together, these results suggested that putative and known gain- and loss-of-function variants produced in these base editing screens may modulate one or multiple hallmarks of T cell function.

### Identification and structural modeling of *PIK3CD* variant hotspots associated with altering T cell polyfunctionality

We next characterized a group of *PIK3CD*-targeting sgRNAs, which enriched across multiple screen readouts and in multiple human donors, thus suggesting that these mutations broadly influence T cell activity (**Fig 3d,e, Extended Data Fig. 3i)**. Class I PI3Ks are obligatory heterodimers composed of a catalytic subunit (p110α, p110β or p110δ) and a regulatory subunit (summarized as p85, with p85a being the most common)(23). PI3Kδ (encoded by *PIK3CD* which forms a heterodimer with p85, encoded by *PIK3R1, 2* or *3*) is predominantly expressed in immune cells and regulates central cell functions such as cell survival and proliferation in response to receptor-tyrosine kinase phosphorylation. Physiologically, p85 recruits and binds to p110δ at the phosphorylated RTKs and stabilizes the catalytic subunit, but inhibits p111 catalytic subunits when occurring as p85 homodimers(24). Activating mutations in the kinase domain of *PIK3CD* (e.g., E1021K) are associated with an immunodeficiency syndrome (Activated PI3K Delta Syndrome, APDS(25,26), which is surprising, given that our screen identified variants that resulted in *improved* polyfunctionality of T cells.

We noted a significant enrichment “hotspot” of multiple base edits in the helical domain of *PIK3CD* resulting in missense mutations in codons 524 (two independent sgRNAs producing Y524C), 525 (two independent sgRNAs producing E525G), 527 (two independent sgRNAs producing E527G), and N529G; additionally, two independent sgRNAs resulting in C416R located in the C2 domain (**Fig. 4a, Extended Data Fig. 6a-c; Extended Data Table 3**). Since the helical domain interacts with the N-terminal SH2 domains (nSH2) and the C2 domain interacts with both nSH2 and the coiled-coil domain (iSH2 domain) of p85, we reasoned that mutations in codons 416 and 524-529 disrupt inhibitory functions of p85, and thereby enable increased catalytic activity of p110δ. In line with this, we also identified *PIK3R1* variants (L570P) and *PIK3R3* (H440R) located in SH2 domains predicted to be important for interacting with the p110 subunits (**Fig. 3d,e**, **Fig. 4b**). *PIK3CD* mutations in residues Y524, E525, D527 and D529 likely interfere with interactions with the opposing Lys/Arg-rich surface of p85, thereby alter the electrostatic interactions at this interface **(Fig. 4b)**. Similarly, the identified mutations in the C416 residue are also adjacent to a Lys (L567 of p85). Thus, mutation to a positive charged amino acid (e.g. C416R) is likely responsible for the functional effect. The L570P mutation identified in *PIK3R1* is likely incompatible with the coiled coil region due to the dihedral angle allowed by the proline substitution, thus disrupting the physiologic interface with p110δ **(Fig. 4b)**.

**Figure 4.**
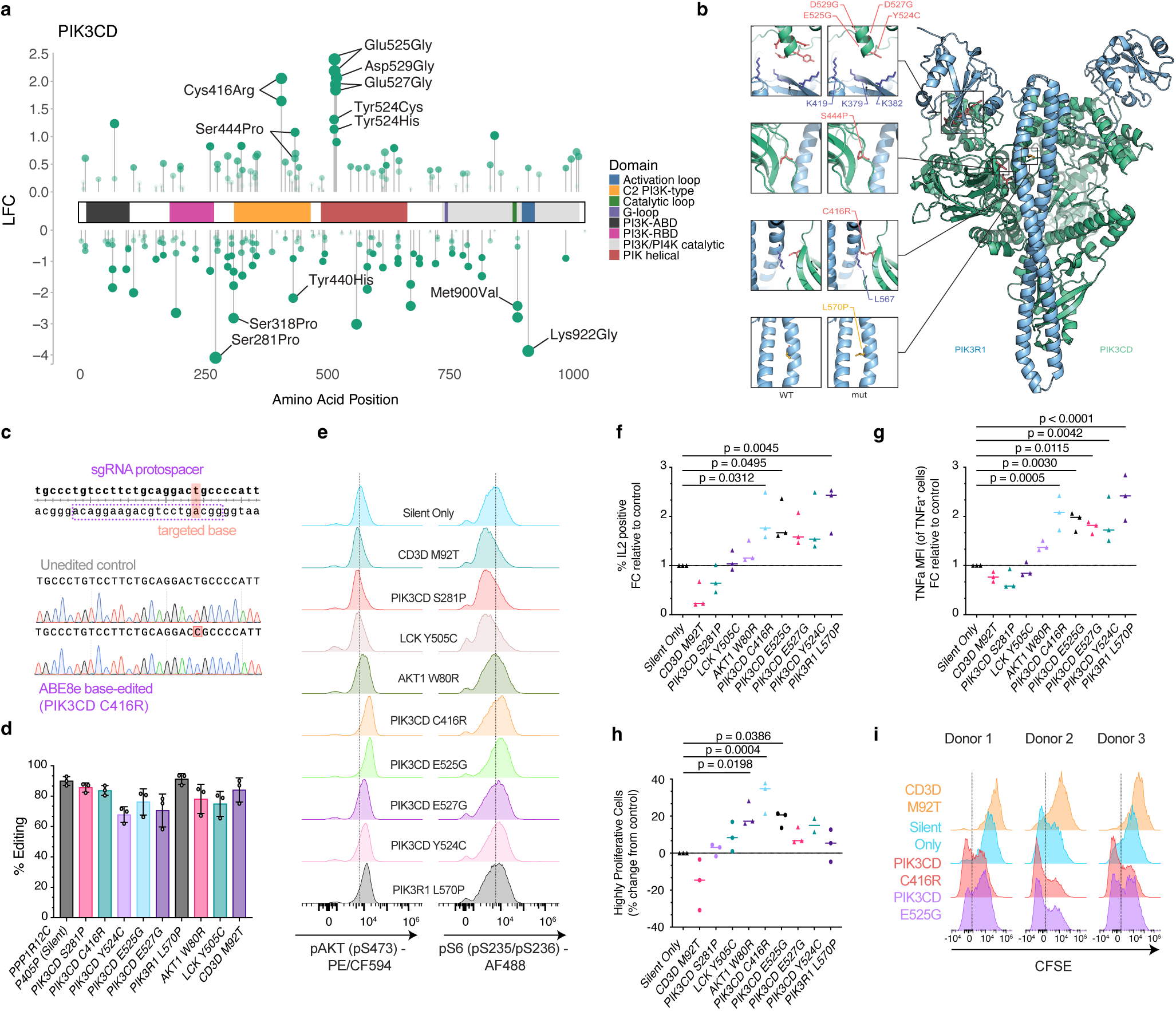
Structure-function analysis of PI3K pathway variants resulting in polyfunctional T cell phenotypes contrasted with LCK mutations predominantly driving isolated T cell proliferation. **a,** Lollipop plot showing LFC of variants produced in base editing screens following repeated stimulation. Indicated are selected variants and their amino acid positions across the gene product of PIK3CD (p110δ) and its highlighted domains. **b,** Predicted structural assembly of PIK3CD (p110δ, green) and PIK3R1 (p85, blue) gene products contributing subunits to active PI3K-δ enzyme. Mutated residues discovered in base editing screens are highlighted (PIK3CD mutations in red, PIK3R1 mutations in yellow). Selected regions of the p110δ/p85 interface are highlighted and contrasted between the wild-type (WT) and mutant (mut) gene products. Mutations Y524C, E525G, D527G, D529G in PIK3CD (green) are predicted to be adjacent to residues K379, K382, K419 (dark blue) in PIK3R1. C416R is adjacent to L567 and L570P localizes to a predicted coiled coil region of PIK3R1. **c,** Representative Sanger sequencing trace from one donor demonstrating adenine base editing of *PIK3CD* generating the C416R mutation compared to unedited control cells from the same donor. sgRNA protospacer and the targeted base are annotated. The targeted adenine (position 4 in the protospacer) is converted to a guanine, leading to a conversion from thymine to cytosine on the antiparallel strand (represented in the sanger traces). **d,** Quantification of base editing efficiency at 10 different loci targeted for validation experiments, as assessed by EditR analysis of Sanger sequencing. *n*=3 independent human donors. **e,** Representative flow cytometry histograms of phosphorylated S6 (pS235/S326) and phosphorylated AKT (pS473) in base edited T cells with indicated mutations following 10 minutes of CD3/CD28 stimulation. Dotted lines represent the approximate median of the histogram in the silent mutation control. Data are representative of one donor; findings were replicated across three independent donors. **f,** Fold change (y axis) of percent IL2-positive T cells with indicated genotypes (x axis) following CD3/CD28 bead stimulation in three independent donors. **g,** Fold change of TNFα mean fluorescence intensity (MFI) in TNFα-positive T cells across indicated genotypes (x axis) following 6-hour CD3/CD28 stimulation. For each donor, the fold change of each variant is normalized to that of the control silent mutation. n=3 human donors. **h,** Proliferation of base edited T cell variants as measured by CFSE dilution following two days of CD3/CD28 stimulation and four days of expansion. Indicated is the frequency (in %) of highly proliferative T cells (CFSE^low^) across indicated genotypes (x axis) in three independent donors. **i,** Representative histograms from flow cytometry experiments in the CFSE experiment (shown in h) for selected mutations in PIK3CD compared to silent controls of deleterious mutation in *CD3D* (M92T). The vertical dotted line indicates the CFSE^low^ gate for highly proliferative cells. One-way ANOVA with Dunnett’s test for multiple comparisons *(panel f, g, h.)*.

### Functional validation of the effect of *PIK3CD* variants on multiple hallmarks of T cell function contrasted with other variants affecting predominantly T cell proliferation

We next sought to functionally characterize several highly enriched *PIK3CD* variants and associated gene variants which were also expected to impact the PI3K axis, including *PIK3R1* L570P as described above, and *AKT1* W80R, an activating mutation downstream of PI3Kδ **(Extended Data Fig. 6d-h)**(21). We reasoned that to determine the specific role of these variants in enhancing *multiple* axes of T cell function, it would be useful to contrast them with variants that apparently only impacted one T cell hallmark in our screens (e.g. proliferation). From both the ClinVar and 12-gene tiling screens, we identified variants of LCK (lymphocyte-specific protein tyrosine kinase) a central regulator of TCR signaling, including Y505C, Y505H and Q506R, all located in the distal C-terminal domain of LCK, and scored strongly in only proliferation assays **(Extended Data Fig. 6i,j)**. Y505C is a ClinVar variant of unknown significance. The tyrosine residue (Y505) is a phosphorylation site that interacts with its own SH2 domain, resulting in an inactive or closed confirmation(27,28) **(Extended Data Fig. 6k)**. Furthermore, as loss-of-function mutation (i.e., negative control for T cell stimulation experiments), we also included a CD3D M92T mutation, a ClinVar VUS which was significantly depleted in multiple readouts in our screens **(Fig 3e, Extended Data Fig. 3k,l)**. To control for non-specific effects of base editor electroporation, we used a sgRNA which only generates a silent mutation in *PPP1R12C* (P405P).

Using the same sgRNA sequences from our screening libraries, we next generated the following single base edits (through simultaneous delivery of BE mRNA and sgRNA via electroporation, see Fig. 1a) in three independent healthy donors: putative *PIK3CD*-activating mutations C416R, Y524C, Y524H, E525G and E527G, and a putative loss-of-function mutation S281P which was depleted in multiple screens **(Fig. 3d-e, Extended Data Fig 3l),** *PIK3R1* L570P, *LCK* Y505C and *AKT1* W80R. We confirmed on-target base editing for these by Sanger sequencing and achieved an estimated average efficiency of 80.25% (ranging from 58 to 93.5%) across all edits and donors (**Fig. 4c,d)**.

We performed a short-term CD3/CD28 stimulation (using crosslinked soluble antibodies) of cells harboring these variants and measured the frequency of S6 and AKT phosphorylation (which are downstream surrogates for PI3K activity). Consistent with our screen readouts, we found increased pS6 (pS235/pS236) and pAKT (pS473) in activating *PI3KCD* variants and *PIK3R1* L570P, and decreased levels in the putative loss-of-function variants *CD3D* M92T and *PIK3CD* S281P (**Fig. 4e)**. Next, we stimulated edited T cells with immobilized OKT3 and soluble CD28, followed by intracellular measurement of cytokines associated with activation (IL2) and effector cytokines (TNFα) using flow cytometry. This resulted in a significantly higher fraction of IL2-positive cells with *PIK3CD* variants C416R and E525G and PIK3R1 L570P (**Fig. 4f**) and increased production of TNFα (**Fig. 4g**) in T cells harboring *PIK3CD* Y524C, E525G and E527G and *PIK3R1* L570P, but not *PIK3CD* S281P or mutations in *AKT1*, *LCK* or *CD3D*. Lastly, we tested the impact of these variants on proliferation rates using the CFSE assay. In all donors, we found increased proliferation in *PIK3CD* C416R, E525G, E527G and Y524C and at a slightly lower rate, *PIK3R1* L570P. As predicted from our screens, *AKT1* W80R and LCK Y505C, but not *PI3KCD* S281P or *CD3D* W92T, also resulted in increased proliferation (**Fig. 4h,i**).

Together, these results allow three important conclusions: first, we validated several variants across different functional axes reflecting the phenotypic readouts used in our screens. Second, we demonstrate that some variants affect multiple T cell functions, while others predominantly influence proliferation. Thus, proliferation as sole readout is insufficient to capture or infer T cell functions critical for effective tumor lysis and emphasizes the importance of the multi-dimensional screen readouts and validation pipelines used in this study. Finally, we also note a spectrum of effect sizes in the signaling, cytokine, and proliferation readouts across the landscape of putative PI3K pathway-activating mutations we validated. This suggests that inferring the effect of a gain-of-function variant based solely on its presence in a signaling cascade is not sufficient to capture the nuance of its influence on tuning signal strength and quality.

### Improving cellular cancer immunotherapies informed by hits nominated in base editing screens

We reasoned that variants that improve multiple axes of T cell activity, such as the selected *PIK3CD* mutations we validated, may be leveraged to improve existing and emerging cell immunotherapies derived from CD8^+^ T cells. To examine this in an epitope-specific manner, we used a previously established co-culture model of the melanoma cell line A375(4), which expresses a common cancer-testis antigen (NY-ESO-1) and human T cells in which we knocked out the endogenous T cell receptor (TCR) and knocked in the cognate NY-ESO-1 T cell receptor (NY-ESO-1 TCR T cells) (**Fig. 5a, Extended Data Fig. 7a-c**). First, we confirmed that we were able to achieve high efficiency base editing in these antigen-specific cells **(Extended Data Fig. 7d)**. Next, we generated several NY-ESO-1 TCR T cells variants, including *PIK3CD* S281P, C416R, and E252G, *AKT1* W80R, *LCK* Y505C, and a silent control mutation in *PPP1R12C* (P405P). A375 cells were plated and co-cultured with NY-ESO-1 TCR T cells harboring the mutants listed above with or without addition of anti-MHC Class I antibody, which blocks TCR-MHC interactions (**Fig. 5a, Extended Data Fig. 7c**).

**Figure 5.**
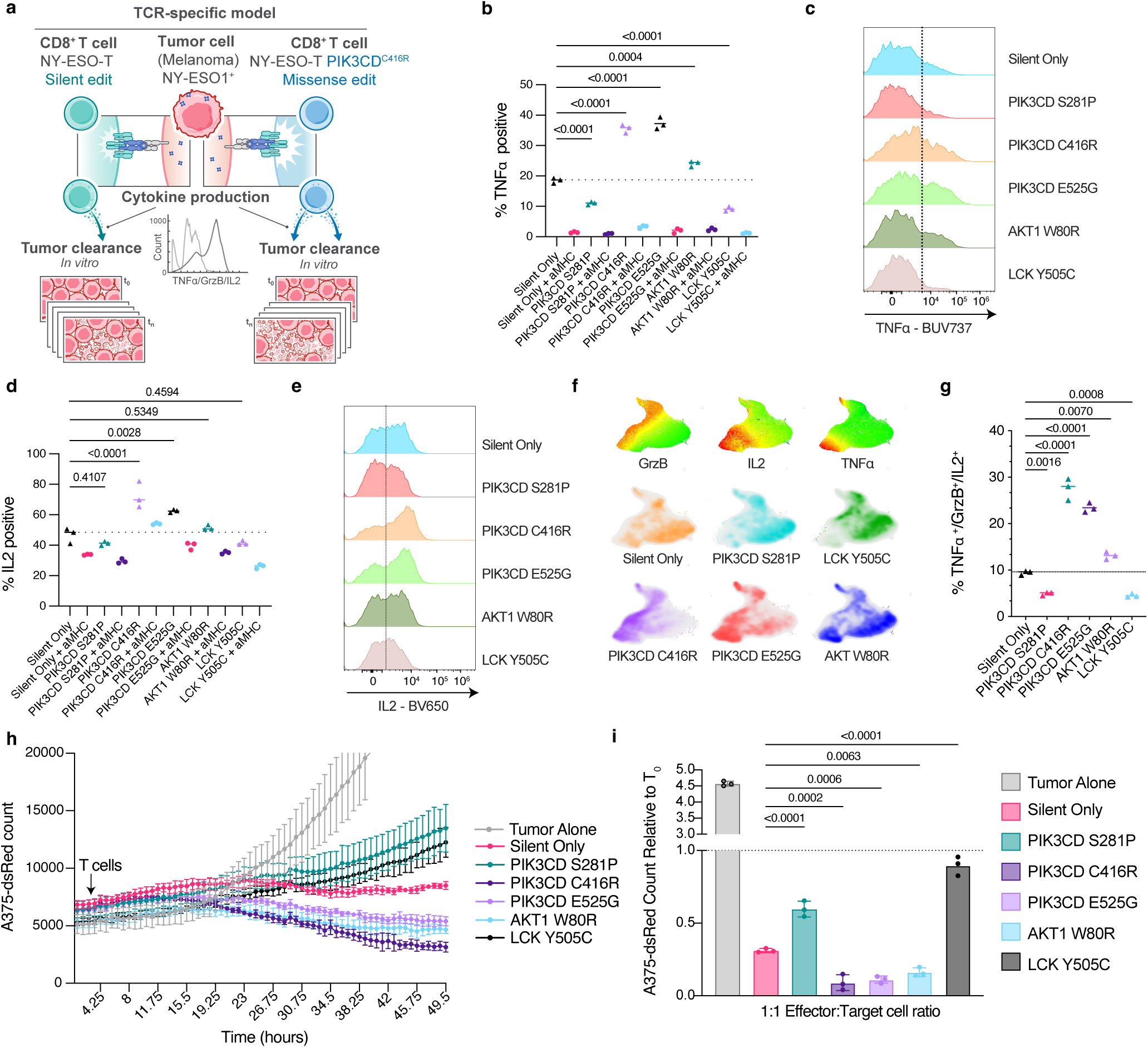
Improving cell-based immunotherapies through base editing of antigen-specific human T cell products. **a,** Schematic representing antigen-matched T cell-melanoma co-culture system leveraged to interrogate variant effects on multiple axes of T cell polyfunctionality and antigen-specific antitumor activity. Primary human T cells engineered to express the NY-ESO-1 T cell receptor (TCR) are co-cultured with NY-ESO-1-presenting A375 melanoma cells. NY-ESO-1 TCR T cells were modified either with a silent mutation or with several mutations identified in base editing screens and predicted to either enhance or diminish T cell polyfunctionality (e.g. *PIK3CD* variants) or predominantly influence isolated T cell functions, such as proliferation (e.g., *AKT1* or *LCK* variants). **b,** TNFα expression in NY-ESO-1-TCR T cells with indicated base edits (x axis) after 8 hours of co-culture with A375, with or without MHC Class I blocking antibody (aMHC) and at an effector to target ratio of 1:1. **c,** Representative flow cytometry histograms of TNFα intensity from variants as in (b). **d,e,** Same as (b) and (d) but showing (d) IL2 positive and (e) representative flow cytometry histograms of NY-ESO-1 specific T cells following co-culture. **f,** Integrated analysis of multiple flow cytometry readouts (using PaCMAP) to show changes on T cell polyfunctionality. Density plots show distribution of cells with indicated variants across multiple measured cytokine readouts from co-culture experiments. Each density plot represents 3 replicates for the indicated variant. **g**, Quantification of T cell polyfunctionality measurement across all flow-cytometry measurements in three independent donors and across indicated tumor-specific T cell genotypes following co-culture with cognate epitope-expressing melanoma cells. **h,** Counts of viable (dsRed-expressing) A375 melanoma cells (y axis) over time (x axis, in hours) plated either alone or co-cultured at an effector-to-target ratio of 1:1 with NY-ESO-1 T cells harboring indicated mutations. T cells were added 4 hours after initial plating, and imaging was performed using an Incucyte instrument at approximately 1-hour intervals. n=3 replicates. **i,** Relative change in cell numbers of A375-dsRed 48 hours after co-culture with NY-ESO-1 variant T cells (effector-to-target ratio 1:1) from an independent experiment. n=3 replicates. One-way ANOVA with Dunnett’s test for multiple comparisons *(panels b, d, g)*.

Following co-culture experiments with or without addition of MHC class I blockade, we isolated T cells and performed intracellular flow cytometry for IL2, TNFα and Granzyme B (GrzB). We found that compared to the silent mutation, the fraction TNFα-producing cells was significantly increased in NY-ESO-1-TCR T cells with *PIK3CD* C416R and E525G, and *AKT1* W80R, and significantly decreased with *PIK3CD* S281P and LCK Y505C **(Fig. 5b,c, Extended Fig. 7e)**. The intensity of TNFα production in *PIK3CD* C416R and E525G T cells was also enhanced, while MHC class I blocking antibody abrogated TNFα production. Similarly, the *PIK3CD* C416R and E525G variants also resulted in increased fractions of both IL2- and GrzB-producing cells **(Fig. 5d,e, Extended Data Fig. 7f)**, as well as higher intensity of GrzB (**Extended Data Fig. 7g,h**). Integrated multi-dimensional analyses across measurements confirmed co-expression of all cytokines in cells with *PIK3CD* C416R and E525G, confirming their polyfunctional state (**Fig. 5f,g, Extended Data Fig. 7i**).

Lastly, we tested the impact of these variants on tumor lysis. We co-cultured NY-ESO-1 TCR T cells with dsRed-expressing A375-melanoma (A375-dsRed, target) cells at 1:1 or 0.5:1 ratio and determined the number of viable tumor cells over time. Compared to the silent mutation, T cells with *PIK3CD* C416R and E525G demonstrated significantly improved tumor killing, while *PIK3CD* S281P or LCK Y505C both demonstrated significantly diminished tumor killing **(Fig. 5h,i, Extended Data Fig. 7j,k)**. Interestingly, T cells with *AKT1* W80R also demonstrated improved tumor lysis. We also noted that across co-culture conditions, the C416R variant showed superior polyfunctionality and tumor-lytic activity compared the other *PIK3CD* variant E525G, suggesting that polyfunctional T cell effects can be tuned through specific base edits. Together, these results demonstrate that synthetic variants detected in base editing screens inform engineering of primary human T cells to make improved cell-based cancer immunotherapies.

## DISCUSSION

The potential clinical efficacy of cell-based cancer immunotherapies, such as TIL transfer products or CAR-T cells, are heavily impacted by the precursor T cell state and function(5–7). Recent studies suggest that point mutations (either germline or engineered) may significantly alter the function of T cells and ensuing cell products. For example, a phospho-silencing mutation in *TSC2* results in antigen-specific, conditional activation and persistence of T cells, and improved anti-tumor activity following ACT(8). Similarly, point mutations in genes encoding for co-stimulatory proteins (e.g., *CD28*) and other signaling molecules (e.g., *PTPN22* or *STAT3*) significantly affect T cell receptor signaling strength, effector and memory phenotype, proliferation, and long-term persistence(9,10), thus, may engender a potentially favorable cell state for production of more efficient cell therapies.

Inspired by such sporadic or naturally occurring examples, we sought to systematically identify variants that may improve the polyfunctionality T cell, thus, enhance the therapeutic activity of products generated from these. While a series of CRISPR-Cas9 studies demonstrated how loss of function screens may identify perturbations that enhance or decrease T cell functions(29–31), these approaches cannot produce specific mutations at sufficient accuracy or scale, and may have several undesirable off-target activities, such as development of aneuploidy(15), a hallmark of cancer. Furthermore, complete gene knockout may result in unknown compensatory upregulation of inhibitory immune checkpoints through loss of steric protein interactions or other mechanisms, and therefore have unexpected immune suppressive consequences(4,13).

In contrast, base editors enable generation of site-specific mutations at single-nucleotide resolution(16) however, application of these methods has been limited due to editing efficiency in primary human T cells and scalability for large-scale discovery(16). Here, we overcame several of these barriers and established methods that enable massively parallel, high efficiency base editing screens in primary human T cells. Additionally, we established optimized procedures for virus- and protein-free single base edits in primary human T cells and engineered T cells that may serve as cell-based immunotherapies.

In multiple base editing screens across different human T cell donors, we simulated acute and tonic T cell receptor engagement, and read out a range of T cell hallmarks that are critical for effective anti-tumor immunity. This multi-modal phenotypic evaluation associated with variants generated in these screens was critical for rational selection of mutations driving a favorable T cell state characterized by activation, proliferation, effector cytokine function and subsequently enhanced tumor lysis. For example, we identified several activating *PIK3CD* mutations that were nominated through the screen and later identified in a series of experiments to confer improved T cell polyfunctionality, proliferation, and improved tumor-lytic capacity, thus, may represent candidate variants that may also improve cell-based immunotherapies. In contrast, other mutations, such as *LCK* Y505C predominantly enhanced cell proliferation, but not cytokine production and ultimately tumor-lytic capacity. Thus, proliferation (or survival) assays alone, which are frequently the readout for large-scale perturbation screens(30,32), are insufficient to nominate potentially useful variants of broad T cell function. Notably, while we identified several activating *PIK3CD* mutations, there were qualitative differences in their phenotypic outputs. For example, C416R consistently produced superior improvement of T cell hallmarks compared to E525G or other activating mutations. Furthermore, mutations in negative regulators of *PIK3CD* gene product p110δ or activating mutations of key nodes downstream of PI3K activity, such as *AKT1* W80R had a significantly lower magnitude in altering T cell polyfunctionality. This suggests that seemingly redundant mutants within the same protein or signaling nodes in the same pathway can have meaningful differential impact on resulting T cell phenotypes. Thus, screens such as the one presented here, may identify opportunities to calibrate and tune T cell functions at a previously underappreciated granularity.

The discovery of activating mutations of *PIK3CD* (and mutations in *PIK3R1/3*) as a means of improving cell-based immunotherapies seems unexpected. Germline mutations in *PIK3CD* have been described to cause a rare syndrome known as activated PI3K delta syndrome (APDS), which is characterized by immunodeficiency and a predisposition for recurrent respiratory infections, yet in a subset associated with auto-immunity(26). This apparent discrepancy – immunodeficiency when occurring in germline and enhanced anti-tumor immunity when introduced in differentiated T cells in our experiments – likely has several explanations. The germline mutations affect immune cells beyond CD8+ T cells, including immunosuppressive T regulatory T cells, and, due to early hyperactivating of CD8+ T cells may be eliminated early during thymic selection, leaving the host deficient for effective cell-mediated immunity. Irrespective, this example highlights an important lesson from these unbiased screens: it is unlikely that one would pick these mutations *a priori* to improve T cell function given the clinical association with immunodeficiency. This emphasizes the power of unbiased discovery coupled with multi-modal functional readouts.

In summary, in this - to our knowledge - first massively parallel base editing screen in primary human T cells, we overcome multiple technical and design barriers, and identified several unexpected mutations that confer enhanced T cell polyfunctionality and antigen-specific tumor killing, thus providing an important blueprint for unbiased discovery to improve existing and future cellular immunotherapies. We also present virus- and protein-free approach for highly efficient base editing, which has several advantages: it is rapid, cost-efficient, and compared to existing approaches for cell-engineering leaves no foreign material after desired base edits. Coupled with the high precision and reduced off-target activities (compared to CRISPR-Cas9 approaches) this approach is safer and will likely also reduce regulatory barriers for such novel cell products to enter clinical testing.

## METHODS

### Primary T cell isolation

Buffy coats from deidentified healthy human donors were obtained from the New York Blood Center. Peripheral blood mononuclear cells (PBMCs) were isolated from buffy coats using SepMate 50 ml conical tubes (StemCell, #85450) containing 15mL of Ficoll-Paque Plus media (1.077g/mL, Cytiva, #17144002) per manufacturer’s protocol. PBMCs were then resuspended in ACK buffer (Gibco, #A1049201) for 5 minutes for red blood cell lysis. CD3+ T cells were isolated from PBMCs via negative selection using the EasySep human T cell isolation kit (StemCell, #17951). All T cell cultures were performed with OpTmizer SFM (Gibco #A1048501) supplemented with 1:40 OpTmizer Supplement (Gibco #A1048501) and 1:100 GlutaMAX (Gibco #35050061) using a humidified incubator at 37 °C with 5% CO_2_ atmosphere.

### Lentiviral production and concentration

HEK-293T cells were passaged at least two times in DMEM (Gibco #11965092) + 10% FBS (Gibco #A5670701), splitting at ∼80% confluence, and then plated at 825,000 cells/well in 6-well plates. 24 hours later, at ∼70-80% confluence, cells were transfected using the TransIT-LT1 system (MirusBio, #MIR2304). Per well, cells were transfected with 500 ng psPAX2 (a gift from Didier Trono, Addgene plasmid #12260), 250 ng VSV-G (a gift from Didier Trono, Addgene plasmid #12259), and 500 ng lentiviral transfer plasmid. 18 hours after transfection, media was replaced with DMEM + 20% FBS. 24 hours after the media change, lentivirus-containing supernatant was harvested from cells and clarified by filtration with a 0.45 uM PES filter. Clarified lentiviral supernatant was concentrated (to 1/10^th^ the original volume) using Lenti-X concentrator (Alstem Bio, #631232) according to the manufacturer’s instructions. Concentrated lentivirus was resuspended in sterile PBS (Gibco, #10010023) and either stored at –80 °C or used immediately for transductions.

### General culture of cell lines

HEK293T and A375 cell lines were cultured in DMEM (Gibco, #11965092) supplemented with 10% FBS (Gibco, #A5670701). All cell lines were cultured at 37 °C in a 5% CO2 humidified incubator. Cells were kept at low passage number and routinely tested for mycoplasma using PlasmoTest (Invivogen, #rep-pt1).

### Lentiviral transduction of primary T cells

Primary human T cells were pre-activated with Dynabeads^TM^ human T-Activator CD3/CD28 beads (Gibco, #11131D) at a 1:1 ratio of cells:beads with 100 IU/ml IL-2 (Chiron #53905-991-01) for 48 hours prior to transduction. T cells were then plated in flat-bottom 96-well plates at 100,000 T cells per well with 1:100 v/v LentiBOOST transduction enhancer (Mayflower Biosciences, # SBPLV10112) and 1:10 v/v of 10x concentrated lentivirus. Plates were centrifuged at 800 x G for 90 minutes at 32 °C in a pre-warmed centrifuge and then cultured overnight at 37 °C. 24 hours after centrifugation, beads were removed using a magnetic rack (Stemcell, #18103), and T cells were expanded in G-Rex 6-well plates (Wilson Wolf, #80240M) with 300 IU/ml IL-2, keeping the density between 0.5-2e6 cells/ml.

### In vitro transcription (IVT) of Base Editor mRNA

Constructs encoding base editors NG-CBE3.9MAX (gift from John Doench and David Root, Addgene #179095) and NG-ABE8e (gift from David Liu, Addgene #138491), and an in vitro transcrition (IVT) template vector (gift from David Liu, Addgene #193843) were obtained from Addgene. Base editors were cloned into the IVT template vector using NEBuilder^®^ HiFi DNA Assembly Cloning Kit (New England Biolabs, #E5520S) (**Extended Data Table 5**) and validated by whole-plasmid sequencing (Plasmidsaurus). The in vitro transcription protocol was adapted from Neugebauer et al(33). The base editor IVT template was simultaneously amplified and poly-T tailed by PCR using primers IVT-F and IVT-R (**Extended Data Table 5**). PCR was performed with Q5^®^ Hot Start High-Fidelity DNA Polymerasewith 25 total cycles (New England Biosciences, # M0493S). The PCR amplicon was purified using a QIAquick PCR Purification Kit (Qiagen #28104), eluted in nuclease-free water (Invitrogen, #10977015), and purity of the amplicon was confirmed by gel electrophoresis prior to IVT. IVT was done using the HiScribe^®^ T7 High-Yield RNA Synthesis Kit (New England Biolabs, #E2040S), with substitution of N-1-methyl-pseudouridine-5’-triphosphate (TriLink Biotechnologies, #N-1081-1) for UTP and co-transcriptional capping with CleanCap^®^ Reagent AG (TriLink Biotechnologies, # N-7113-1). For large-scale mRNA production, 320ul of IVT reaction was performed for each editor construct. IVT mRNA was then precipitated by mixing with 0.5 v/v Lithium Chloride (Invitrogen, #AM9480) and incubating at −20 °C for 30 minutes. mRNA was pelleted by centrifugation at 15,000xG for 20 minutes at 4 °C, washed once with ice-cold 70% ethanol, and then supernatant was removed and pellet air-dried at room temperature for 5 minutes. The mRNA pellet was resuspended by gentle pipetting in 500 µl Ambion THE RNA Storage Solution (Thermo Fisher, #AM7000) per 320 µl of initial IVT reaction. mRNA concentration was quantified using a Nanodrop Spectrophotometer (Thermo Fisher), and mRNA was further diluted to a final concentration of 1.5 µg/µl. Appropriate purity and size of the mRNA products was confirmed using an RNA tapestation kit (Agilent, #5067-5579). Prior to use in screens or validation experiments, editing activity of the BE encoded by the mRNA was confirmed by knockout of a constitutively expressed T cell protein with a validated high efficiency sgRNA.

### Design of base editor sgRNAs for targeted gene disruption experiments

For targeted gene knockout experiments to validate base editor efficiency, sgRNAs creating mutations in *CD2* splice sites were generated using SpliceR (https://github.com/MoriarityLab/SpliceR)(18). sgRNAs creating premature stop codons in *B2M* were generated using iSTOP (https://github.com/CicciaLab/iSTOP)(34). sgRNAs ablating the start codon of *B2M*were generated using the Base Editor Design Tool (https://github.com/mhegde/base-editor-design-tool). All sgRNAs were ordered as full-length synthetic sgRNAs with modified bases (2’ O-methyl analog on first and last 3 bases; 3’ phosphorothioate between first 3 and last 2 bases) (Synthego). A full list of sgRNAs used for single-guide perturbation experiments are included in **Extended Data Table 6**.

### Base editing of primary T cells

For single and multi-plex (non-library) base-editing, T cells were activated for 48h in 100 IU/ml IL2 with a 1:1 ratio of CD3/CD28 activator beads, as indicated. T cells were then washed 1x with PBS, resuspended at 1e6 / 20 µl in P3 nucleofection buffer (Lonza, #V4XP-3032), and 1e6 cells were combined with 4.5 µg BE mRNA and 100 pmol sgRNA (Synthego). T cells were electroporated at 1e6 per cuvette well in a Lonza 4D nucleofector using electroporation program EO-115, and then 100 µl pre-warmed media was immediately added to each cuvette well. Cells were recovered in the cuvette for 15 minutes at 37 °C, and then cultured in OpTmizer SFM at 1e6/ml with 300 IU/ml IL2. Editing efficiency was assessed at least 4 days after nucleofection by flow cytometry or Sanger sequencing (Azenta).

### Lentiviral-based single-guide editing of primary T cells

For single-guide perturbation pilot experiments using lentiviral guide integration, sgRNAs were cloned into a modified CROPseq-mTurquoise-PuroR backbone (CropSeq-mTurq) using the GeCKO single-guide cloning protocol. Briefly, for each sgRNA, two oligos were ordered, Oligo1: 5’-CACCG[20nt guide]-3’, Oligo2: 3’-C[revcom of 20nt guide]CAAA-5’. The oligos were annealed and phosphorylated, and then ligated into BsmBI-digested, dephosphorylated CROPseq backbone with T4 ligase. A full list of cloning oligos is included in **Extended Data Table 7**. Ligation products were transformed into Stbl3 bacteria (Thermo Fisher, #C737303), DNA was purified from bacterial cultures using a HiSpeed Plasmid MidiPrep Kit (Qiagen, #12643), and cloned products were validated with whole-plasmid sequencing or Sanger sequencing of the insert. CropSeq lentivirus was generated, and primary T cells were transduced with lentivirus as described above. After lentiviral transduction cells were left to recover for 72 hours prior to nucleofection. For Cas9-editing proof-of-concept experiments, Cas9 was then introduced using nucleofection of CleanCap 5moU Cas9 mRNA (Trilink, #L-7206). For base-editing experiments, base editors were introduced using IVT mRNA encoding either NG-ABE8e or NG-CBE3.9MAX. To this end T cells were washed once in PBS and resuspended in complete P3 nucleofection buffer (Lonza, #V4XP-3032). 1e6 cells were then mixed with either Cas9 or base editor mRNA (4 - 4.5 µg) and nucleofected with program EO-115 using a 4D Nucleofector (Lonza). Immediately after nucleofection, 100 µl complete T cell media was added and cells were left to recover for 15 min at 37 °C before transfer to culture plates at 1e6/ml with T cell media supplemented with 300 IU/ml IL2. Editing efficiency was assessed 4-5 days after nucleofection using a Cytek Aurora flow cytometer (Cytek). Transduced cells were identified by gating on viable, mTurqoise positive cells and target gene KO was assessed by comparing target gene expression in mTurqoise positive vs. mTurquoise negative cells and cells which had not received Cas9 mRNA or Protein.

### Flow cytometry and flow cytometry assisted cell sorting (FACS)

#### Surface staining

For all surface staining experiments, cells were collected, washed 1x with ice-cold FACS buffer, and stained on ice for 20 minutes in FACS buffer with surface antibodies at dilutions pre-determined by titration experiments. When multiple brilliant dyes were used together, Brilliant Stain Buffer Plus (BD, #566385) was added according to the manufacturer’s instructions. Cells were then stained with Sytox Green Ready Flow Reagent (Invitrogen, #R37168) according to the manufacturer’s protocol, and analyzed on a Cytek Aurora flow cytometer (Cytek). In cases where staining with Sytox Green was not feasible (fixed samples, fluorophore overlap), cells were instead stained with a 1:1000 dilution of Zombie NIR Viability Dye (Biolegend, #423105) in PBS on ice for 10 minutes prior to surface antibody staining.

#### Intracellular cytokine staining

For intracellular cytokine staining experiments, cells were collected and stained on ice for 10 minutes with a 1:1000 dilution of Zombie NIR in PBS. Cells were then stained for surface markers of interest for 20 minutes on ice. Surface-stained cells were fixed and permeabilized using the eBioscience FoxP3 / Transcription Factor Staining Buffer Set (Invitrogen, #00-5523-00), stained for intracellular cytokines of interest and analyzed on a Cytek Aurora flow cytometer. When multiple brilliant dyes were used together, Brilliant Stain Buffer Plus (BD, #566385) was added according to the manufacturer’s instructions.

#### Intracellular phosphoprotein staining

For all phospho-flow cytometry experiments, cells were stimulated at 37 °C for a pre-determined time period (additional information in “Phospho-flow cytometry of T cell variants” section) Following stimulation at 37 °C, cells were immediately fixed by adding an equal volume of pre-warmed BD Cytofix Buffer (BD, #554655), permeabilized with BD Phosflow Perm Buffer III (BD, #558050), and then stained with antibodies against intracellular phospho-targets for 30 minutes at room temperature in the dark. Samples were run on a Cytek Aurora cytometer and analyzed using Flowjo v10.

### High-dimensional visualization of flow cytometry data

For high-dimensional visualization of intracellular T cell cytokine production, samples were indexed based on T cell variant, concatenated, and visualized using PaCMAP dimensionality reduction using all default parameters(35). Dimensionality reduction was performed based on four intracellular proteins: IFNγ, IL2, TNFα, GrzB. All analysis was and visualizations were performed in FlowJo V10.8.1 with the PaCMAP plugin.

### Design of base editor libraries

For the ClinVar library, the KEGG gene set “T CELL RECEPTOR SIGNALING PATHWAY” (hsa04660) was used to generate a starting list of 108 target genes. Genes encoding secreted proteins (interleukins, *IFNG*) were excluded, as the impact of their perturbation on the cell of origin is difficult to determine. Using the final list of 102 genes (**Extended Data Table 1**), we used the Base Editor Design Tool (https://github.com/mhegde/base-editor-design-tool) to design a library tiling each gene. The experimental perturbation library was filtered to include only sgRNAs introducing missense mutations in amino acids with known variants classified as “variant of unknown significance,” “likely pathogenic,” or “pathogenic.” To broaden our mutation library, we did not filter sgRNAs making non-identical missense mutations at these loci (e.g. an sgRNA that makes a Y100H missense mutation at Tyr100, when the known ClinVar variant is Y100C, would not be filtered). sgRNAs containing a “TTTT” sequence or BsmBI cut site were also excluded. The negative control library consisted of 300 randomly-selected guides predicted to make only silent mutations in these genes, and 300 guides with no target base in the 5-nucleotide editing window (“empty window”). 200 additional control guides were selected which tiled the *PPP1R12C* gene, which is not expected to play a role in T cell function. For positive controls (i.e. expected dropouts) 50 genes classified as pan-species essential by the Bayesian Analysis of Gene Essentiality 2 (BAGEL2)(36) were selected as targets. We generated a library of 600 sgRNAs predicted to perturb these genes: 300 sgRNAs introducing splice site mutations, including both splice acceptor and splice donor sites, and 300 sgRNAs introducing proline mutations (i.e. mutation of any non-proline amino acid to proline), which are expected to disrupt protein structure.

For the 12-Gene tiling library, target genes were selected based on their central roles in orchestrating T cell responses to extracellular cues. Each gene was tiled with sgRNAs using the Base Editor Design Tool as above. Filtration was performed to remove sgRNAs containing a BsmBI cut site. For negative controls, 300 guides introducing silent mutations and 300 empty-window guides targeting these genes were included, in addition to 200 *PPP1R12C* tiling guides, as described above. For positive controls, the identical library of 600 sgRNAs was used as described in the ClinVar library.

### Visualization of targeted geneset for ClinVar base editor library

To visualize known functional protein interactions and groups of targets included in the ClinVar library the gene list was uploaded to the STRING database (https://string-db.org/) and interactions were visualized using the full STRING network including confident interactions based on textmining, experiments, databases, co-expresison, neighborhood, gene fusion and co-occurrence using the highest confidence setting (0.9)(37). The protein interactions were then clustered using STRING and k-means clustering with the number of clusters set to 8. The resulting clusters were annotated based on biological function.

### Base editor library cloning

Custom oligonucleotide pools (TWIST Bioscience) encoding the individual sgRNA sequences and flanking regions for targeted subpool amplification and cloning were used for pooled golden-gate cloning of sgRNA gene editing libraries into the CropSeq vector as previously described(38). The following sequence was used for all oligonucleotide pools: 5′-[Forward Primer] CGTCTCA***CACC***G [sgRNA, 20 nt] ***GTTT***CGAGACG [Reverse Primer]-3’. The individual library pools were amplified using subpool specific primers and KAPA HiFi HotStart ReadyMix (Roche, #KK2601). After PCR, the reactions were cleaned using the QIAquick PCR purification kit (Qiagen, #28104) according to manufacturer instruction and elute in 50 μL TE buffer (Invitrogen, #12090015). To prepare the CropSeq-mTurq backbone for pooled golden-gate cloning, 5 µg of sequencing verified plasmid were restriction digested using BsmBI V2 (NEB, #R0739S) in a 50 µl reaction with NEB3.1 buffer (NEB, #B7203S) for 2 hours at 55 °C. After digestion the linearized backbone was dephosphorylated by addition of 2 µl rSAP (NEB, #M0371S) and incubation at 37 °C for 1 hour followed by heat inactivation of the enzymes at 80 °C for 20 minutes. The reaction was then loaded on a 1% agarose gel confirming the release of a ∼1.8 kb fragment. The linearized backbone was extracted from the gel using the Zymogen Gel DNA recovery Kit (#D4001) followed by additional clean up using 1x SPRI bead selection (Beckman Coulter, #B23318) and eluted in TE buffer. Next, the linearized backbone and amplified oligonucleotide pool were mixed at equimolar ratio and assembled using a 50 µl golden-gate reaction containing 1x Tango Buffer (Thermo Fisher, #BY5), 1 mM DTT (Thermo Fisher, #R0861), 1 mM ATP (NEB, # P0756S), 1 µl Esp3l (Thermo Scientific, #ER0451), T7 ligase (NEB, #M0318S) and nuclease free water. The reaction was conducted in a thermocycler using incubation at 37 °C for 5 minutes followed by 20 °C for 5 minutes for a total of 99 cycles (final holding stage at 4 °C). After the reaction had been completed the final product was cleaned using 1x SPRI beads and eluted for 1 minute at room temperature in 11 µl TE. Next, 10 µl of the reaction were electroporated in 50 µl Lucigen Endura electro competent cells (Lucigen, #602421) using 1 mm electroporation cuvettes (Biorad, #1652089) and a BioRad Gene Pulser Xcell (Biorad, #1652660) set to 10 µF, 600 Ohms, 1.8 kV. Immediately after electroporation 1 ml of prewarmed recovery medium (Lucigen, #800261 was added to the bacteria and the cells were incubated for 1 hour at 37 °C shaking at 250 rpm in a bacterial incubator. Thereafter the cells were plated on a 25×25 cm LB-agar plate with 100 µg/ml Carbenicillin and incubated for 14 hours at 32 °C. Transformation efficiency and library coverage was in parallel monitored by serial dilution(39). To extract the final plasmid libraries the bacteria were scraped from the LB-agar plates into LB buffer and collected by centrifugation. The plasmid DNA was extracted using Qiagen HiSpeed Maxi kit (Qiagen, #122663) with one column per 2-gram bacterial pellet. The final libraries were quantified using a NanoDrop and the library distribution was assessed using next generation sequencing (see below) before preparation of lentivirus as described above.

### Base editor screening

#### Massively parallel base editing of primary T cells

For each screen, buffy coats from two healthy human donors were purchased. Primary CD3 T cells were isolated, bead-activated, and transduced with concentrated library-containing lentivirus as described above. At least 20e6 T cells per donor were transduced to ensure appropriate (>500x cells per guide) library representation. While on activator beads, cells were cultured in 100 IU/ml IL2, and following bead removal, cells were cultured in 300 IU/ml IL2. 72 hours after lentiviral spinoculation, T cells were resuspended in P3 buffer at 1e6 cells / 20 µl. Per nucleocuvette well, 1e6 T cells were electroporated with 4.5 µg ABE8e mRNA in a Lonza 4D nucleofector using the electroporation program EO-115. At least 24e6 T cells per donor were electroporated to maintain library representation. Following electroporation, cells were immediately recovered in 100 µl pre-warmed media at 37 °C for 15 minutes, and then 24-32e6 T cells per donor were cultured in 40 ml OpTmizer SFM in G-REX 6-well plates. Media and IL2 were refreshed every 48-72 hours for the duration of culture. For downstream functional/phenotyping assessment and sorting, cells were allowed to edit for at least 7 days after electroporation prior to assay initiation. Individual downstream assays are described in separate subsections.

#### Long-term T cell expansion

For long-term expansion of library-edited T cells, 20e6 cells were restimulated every ∼7-14 days with CD3/CD28 activator beads at a 1:1 bead:cell ratio for 48 hours, followed by bead removal and G-REX expansion. Media and cytokines were refreshed every 72 hours by half-media exchange.

#### Cytokine production sort

8 days after ABE electroporation, library-edited T cells were collected and resuspended in OpTmizer without IL2 for 24 hours. Then, T cells were activated for 6 hours in T175 flasks pre-coated with 1 µg/ml anti-CD3 (OKT3) (Miltenyi #130-093-387), with addition of 2 µg/ml soluble anti-CD28 (Miltenyi, #130-122-350) and eBioscience 1x Protein Transport Inhibitor Cocktail (Thermo Fisher, #00-4980-93). T cells were then collected for intracellular cytokine staining. Fixation and permeabilization was performed using the eBioscience FoxP3/Transcription Factor Staining Buffer set as described above. Sorting was performed on a Sony MA-900.

#### CFSE proliferation sort

6 days after ABE electroporation, T cells were stained at 20e6/ml in PBS with 1 µM CFSE (BD Biosciences, #) in a 37 °C water bath for 15 minutes. An aliquot of cells was analyzed by flow cytometry to confirm brightness and uniformity of CFSE staining. CFSE-stained T cells were resuspended at 1e6/ml and Dynabeads^TM^ human T-Activator CD3/CD28 beads (Gibco, #11131D) were added at a ratio of 1:10 beads:cells. Cells were activated on beads for 48 hours, followed by bead removal, and then allowed to proliferate for 4 additional days before collection, flow cytometric sorting, and NGS.

#### CD25 activation sort

12 days after ABE electroporation, library-edited T cells were collected, stained with Zombie NIR viability dye, and then stained for CD25 and CD8 on ice for 20 minutes prior to sorting.

#### DNA extraction and next-generation sequencing

For unfixed samples (long-term proliferation, CD25, and CFSE screen readouts), DNA was extracted from library-edited T cells using a DNeasy Blood & Tissue Kit (Qiagen, #69504) according to the manufacturer’s protocols with buffer AL, and eluted in 100 µl buffer EB (Qiagen, #19086) per 5e6 T cells. For fixed samples (cytokine screen readout), DNA was extracted as above with the following modification: instead of incubating at 56 °C for 10 minutes in Buffer AL and proteinase K, samples were incubated overnight at 65 °C in buffer AL and proteinase K with shaking at 1000 RPM. Sample amplification and indexing PCR was performed with ExTaq polymerase (Takara Bio, ## RR001A) under the following conditions: for unfixed and unsorted samples, 10 µg gDNA input per 100 µl PCR with 24 cycles. For unfixed sorted samples, 3 µg gDNA input per 100 µl PCR with 24 cycles. For fixed samples, 1µg gDNA input per 100 µl PCR with 26 cycles. Each gDNA sample was split across multiple PCRs (up to 8 parallel PCR per sample) and then PCR products were pooled to minimize skewing of guide distribution induced by PCR amplification. Pooled PCR products for each sample were cleaned up with a 1x SPRI, quantified by D5000 Tapestation, and then pooled targeting 500x read coverage per sgRNA in each library and sequenced with 75 bp Read 1, 8 bp Index 1 on a NextSeq 550 sequencer (Illumina). Primers used for sample indexing PCR are listed in **Extended Data Table 5**.

### Bulk screen analysis

Fastq files were trimmed at the 5’ end using cutadapt to remove the constant sequences. Analysis of guide enrichment was performed using MAGeCK(40). The MAGeCK ‘count’ command was used to determine the number of reads for each guide. Next, the MAGeCK ‘test’ command was used to determine enriched and depleted guides in sample versus control conditions. MAGeCK’s robust rank aggregation method was used to determine statistical significance for each comparison. For all long-term proliferation analyses in both ClinVar and 12-gene tiling screens, sgRNA distribution in the edited T cell population at the indicated timepoint was compared to the sgRNA distribution in T cells which were transduced with the sgRNA library but not electroporated with base editor (labeled as “Day 0 post-electroporation” in timecourse plots), thus providing a baseline sgRNA distribution in cells.

Analysis and visualization were conducted in R using the MAGeCK outputs. For guides associated with multiple edit types in the 12-gene tiling screen (e.g. missense, silent), the guide was classified according to the anticipated most-impactful edit (i.e. splice donor>splice acceptor>missense>utr>silent) for lollipop plot visualization. For the density plot analysis, a one-sided 5% threshold was established based on the distribution of empty window and silent mutations pooled together. For the analysis of CD3 complex and essential gene guides across donors, a linear model was fit to the data to determine R^2^ values. Density plots, scatterplots, lollipop plots, and volcano plots were visualized using the ggplot2 package.

### Generation, visualization, and analysis of structures protein variants from base editor screens

AlphaFold2-predicted structure of the PIK3CD-PIK3R1 complex was retrieved from Model Archive (DOI: 10.5452/ma-t3vr3)(41). Gene names AKT1 and LCK were mapped to UniProt identifiers P31749, P06239, respectively, using the UniProt ID Mapping service(42). UniProt IDs were queried against the AlphaFold Database (https://alphafold.ebi.ac.uk) to retrieve the corresponding AlphaFold2-predicted structures, AF-P31749-F1-model_v4, AF-P06239-F1-model_v4, respectively. All structures were visualized in PyMOL v2.5.7 corresponding scripts are available at https://github.com/gnikolenyi/izar_vis.

### Sanger sequencing validation of single base-edits

Following sgRNA and base editor electroporation, T cells were cultured for at least 4 days to allow editing to occur. Then, 5e6 cells were collected, and DNA was extracted using a DNeasy Blood and Tissue kit (Qiagen, #69504) according to the manufacturer’s instructions. Unique PCR amplification primers were designed to amplify a ∼400nt region centered around the base-edited locus using PrimerBlast (NCBI). The region was amplified using NEB Q5 HotStart HiFi (NEB, #M0494S). After initial denaturation at 98 °C for 30 seconds the targets were amplified by 35 cycles of 10 seconds denaturing at 98 °C, 30 seconds annealing at 67-69 °C (primer specific) and 30 seconds extension at 72 °C, followed by a final extension for 2 minutes at 72 °C and a infinite cooling step at 4 °C. After the PCR the product was gel-extracted, purified using a ZymoClean Gel DNA Recovery Kit (Zymogen, #D4001), further purified with a 1x SPRI selection (BD, #B23318), and Sanger sequenced (Azenta). Editing efficiency of the amplicon was assessed using EditR(43). A complete list of all primers used for sanger sequencing is included in **Extended Data Table 8**.

### Assessment of cytokine production by T cell variants

Assessment of cytokine production by T cell variants from screens was performed as in the screen cytokine sort with the following modifications: T cells were stimulated for 6 hours in 96-well plates instead of flasks, at a concentration of 100,000 T cells per well.

### Assessment of T cell variant proliferation by CFSE

Assessment of proliferation by T cell variants was performed as in the screen CFSE sort with the following modifications: CFSE-stained T cells were stimulated for 2 days with 1 µg/ml plate-coated OKT3 (Miltenyi, #130-093-387) at 100,000 cells/well in 96-well plates, followed by passaging to a fresh plate and expansion for an additional 4 days prior to flow analysis on a Cytek Aurora cytometer.

### Phospho-flow cytometry of T cell variants

Prior to downstream phospho-flow cytometry analysis, T cells were first rested from exogenous stimulation for 24 hours. To assess phosphorylation of key intracellular targets, T cells were stained on ice with 20 µg/mL anti-CD3 (BD, #555329) and 20 µg/mL anti-CD28 (BD, #555725) solution for 20 minutes followed by 20 minutes of antibody cross linking on ice with 2 µg/mL goat-anti mouse solution (BD, #53998). T cells were then activated via an incubation period at 37 °C in FACS buffer (PBS with 2% FBS and 2 mM EDTA) for which the timing was optimized for each intracellular target. T cells were then immediately fixed, permeabilized, stained, and analyzed by flow cytometry as described above.

### Generation of cancer lines for in vitro and in vivo experiments

A375-dsRed-NLS cells expressing dsRed with a nuclear localisation sequence to facilitate automated cell counting were generated previously described(38).

### CRISPR-Cas9 editing of primary T cells

Single-guide CRISPR-Cas9 ribonucleoprotein (RNP) perturbation of primary T cells were performed as previously described(4). Briefly, virus-free knock out (KO) of target genes in primary human T cells were generated by Cas9 ribonucleoprotein (RNP) electroporation using a 4D Nucleofector (Lonza) and P3 primary cell nucleofection reagents (Lonza, #V4XP-3032). Target sequences for gene KO were identified using CRISPick(44) (**Extended Data Table 6**). CRISPR RNA (crRNA; IDT) and transactivating CRISPR RNA (tracrRNA; IDT) were mixed at equimolar ratio in nuclease free duplex buffer (IDT) and incubated for 5 min at 95 °C followed by cooling to room temperature to assemble the target specific guide RNA (gRNA). The gRNA was then mixed with recombinant Cas9 protein (MacroLab) at 1:10 molar ratio and incubated for 15 min at 37 °C. The final RNP was stored on ice until use. To prepare the cells for nucleofection, T cells were collected and washed once with PBS. 1e6 T cells were resuspended in 20 µl P3 electroporation buffer with P3 supplement (Lonza. #V4XP-3032) as per manufacturer instructions and combined with 3 µl RNP. Cells were electroporated using program EH115. Immediately after, 100 µl T cell media was added to the cells and the cells were allowed to rest for 15 min at 37 °C before transfer to culture plates with T cell media supplemented with 300IU/ml IL2. Edited T cells were cultured for at least 3 days prior to assessment of KO editing efficiency by flow cytometry.

### Generation of NY-ESO-1 T cells

Polyclonal T cells were isolated from PBMCs (see above) and pre-activated with Dynabeads^TM^ human T-Activator CD3/CD28 beads (Gibco, #11131D) at a 1:1 ratio of cells:beads with 100 IU/ml IL-2 (Chiron). After 72 hours cells were removed from beads and T cells were engineered to express NY-ESO-11 (1G4) specific T cell receptor through orthotopic knock-in at the TRAC locus as previously described(45,46). The homology directed repair (HDR) template was generated by PCR from a template plasmid (a gift from Alexander Marson, Addgene plasmid #112021)(45) using primers containing truncated Cas9 target sequences (tCTSs). This is to facilitate transport of the template to the nucleus via interaction with the Cas9 RNP complex containing a nuclear localization signal and increases editing efficiency(45). After PCR, the product was purified using SPRI selection and quantified using Tapestation D5000 reagents. Cas9-RNP targeting *TRAC* were assembled as described above and the final RNP were incubated with the HDR template for 5 minutes at room temperature. The precomplexed HDR-RNP complex was electroporated into primary human T cells as described above using P3 buffer (Lonza). The T cells were cultured in T cell media with 300 IU/ml IL-2 for 7-10 days. On day 7-10, cells were stained with anti-CD8 and NY-ESO-11 TCR specific dextramer (Immudex, # WB3247-PE). Dextramer positive CD8 T cells were sorted using a Sony MA-900 Cell Sorter. After sorting, the T cells were cultured in T cell media with 300 IU/ml IL-2, further expanded, frozen in aliquots, and stored in liquid nitrogen storage until further use.

To expand NY-ESO-1 specific T cells for experiments we adopted a rapid expansion protocol as previously described(4,38). Edited NY-ESO-11 specific T cells were thawed and rested for 2-3 days in standard culture conditions. To expand the cells, 1e6 T cells were seeded with 100e6 allogenic pooled PBMC irradiated in a GREX-10 bottle with 40 mL of REP media containing (50% AIM V (Thermo Fisher, #12055091) and 50% T cell media) supplemented with 30 ng/mL of anti-CD3 (OKT3, Miltenyi, #130-093-387) and 3000 IU/mL IL-2. On day 2, an additional 3000 IU/ml IL2 was added. On day 5, fresh media and IL2 were added by half media exchange. On day 7, cells were counted. If the expansion exceeded >6e7 cells, cells were reseeded in individual wells of a 6-well GREX plate with 2e7 cells per well using AIM V media; If expansion remained <6e7 cells the cells remained in the GREX-10 bottle. In either condition 3000 IU/ml IL2 were added. On day 10 and 12, fresh media and IL2 were again added by half media exchange. On day 14, the expansion was completed, and cells were cryopreserved and stored in liquid nitrogen until use in experiments.

### Melanoma:NY-ESO-1 T cell co-cultures

To assess antigen specific killing of melanoma target cells by base edited cytotoxic CD8 T cells we cocultured human A375 melanoma cells with endogenous expression of the cancer testis antigen NY-ESO-11 with NY-ESO-11 TCR engineered CD8 T cells as previously described(4). A375 target cells expressing dsRed-NLS for automated counting of target cell nuclei were seeded at 10e4 cells per well in black walled, clear bottom 96 well plates (Corning, #3603). Target cells were seeded in 100 µl DMEM per well supplemented with 10% FBS. To monitor apoptosis, a green-fluorescent caspase3/7 activity reporter dye (CellEvent; ThermoScientific, #C10423) was added (2 µM final concentration in coculture). For MHC-I blocking control conditions, 50 µg/ml anti-HLA-A,B,C (W6/32; Invitrogen, #MA1-19027) were added. After 4 hours the plates were imaged on a Celigo Imaging Cytometer (Nexelcom) for a t_0_ count of viable target cells. Next, NY-ESO-11 specific T cells were washed once in complete T cell media, resuspended, counted and added in T cell media without IL2 with increasing effector:target ratio as indicated. The cocultures were incubated for 48 hours in a humidified incubator (37 °C, 5% CO_2_) equipped with an Incucyte (Sartorius) for continuous monitoring of cell proliferation and target cell killing. At the 24 and 48 hours timepoint the plates were also imaged on a Celigo Imaging Cytometer (Nexelcom). To quantify target cell killing, melanoma cells were counted using the nuclear dsRed signal identified by the Celigo Imaging Cytometry software and final counts were normalized to matched t_0_ counts. To monitor continuous target cell killing and proliferation the melanoma cell dsRed intensity were measured over the same three regions in each well every 45 minutes using Incucyte’s Cell-by-cell analysis software.

### Flow cytometry for antigen-specific T cell activation and cytokine production

To assess antigen-specific activation and induction of cytokine production, NY-ESO-1 TCR T cells were co-cultured with A375 melanoma target cells at 1:1 effector:target ratio in 96-well plates in triplicate (10,000 T cells and 10,000 melanoma cells per well). At co-culture initiation, 1x Protein Transport Inhibitor Cocktail (Invitrogen, #00-4980-93) was added to facilitate intracellular cytokine accumulation. For MHC-I blocking control conditions, 50 µg/ml anti-HLA-A,B,C (W6/32; Invitrogen, #MA1-19027) were added. 8 hours after co-culture initiation, T cells were harvested, stained with Zombie NIR fixable viability dye (BioLegend, 423105) at a dilution of 1:1000, and then stained for surface markers of interest. Next, T cells were fixed and permeabilized using the eBioscience FoxP3/Transcription Factor Staining Buffer Set (Invitrogen, #00-5523-00) following manufacturer’s instructions and stained for intracellular cytokines for 30 minutes at room temperature. Samples were run on a Cytek Aurora flow cytometer.

### Statistical Analysis

For visualization of Volcano plot data, the FDR cutoff for positive and negative hits was set to FDR<0.05. For all validation experiments, one-way ANOVA with Dunnett’s test was used to compare each experimental variant to the silent mutation-only control. Statistics for all screen validation experiments were performed in Prism 10 (GraphPad).

## ACKNOLWEDGEMENTS

B.I. is supported by National Institute of Health grants, R37CA258829, R01CA280414, R01CA266446, U54CA274506, and additional by the Pershing Square Sohn Cancer Research Alliance Award, the Burroughs Wellcome Fund Career Award for Medical Scientists; a Tara Miller Melanoma Research Alliance Young Investigator Award; the Louis V. Gerstner, Jr. Scholars Program; and the V Foundation Scholars Award. Medical illustrations were prepared by Dr. Uta Mackensen.

## AUTHOR CONTRIBUTIONS

B.I. and Z.H.W. conceived of the study. B.I. provided overall supervision with support from J.C.M. Z.H.W., P.S. and J.C.M. planned, designed, and executed all key experiments. S.B.S., G.L., M.M., P.H. and S.A. performed experiments. N.K. performed computational analyses of screens with support from Z.H.W. and Z.D.D. G.N. performed structural modeling and visualizations. N.V., M.A., J.D.M. and A.D. provided additional guidance for the design, execution and interpretation of screens. Z.H.W., P.S., J.C.M. and B.I. wrote the manuscript, with input and approval from all authors.

## DATA AVAILABILITY

All screening data results are provided as supplementary tables to the manuscript.

## CODE AVAILABILITY

Code for generating *in silico* predicted structures is deposited here: https://github.com/gnikolenyi/izar_vis

## EXTENDED DATA FIGURE LEGENDS

**Extended Data Figure 1. Optimization of workflows for base editing in primary human T cells. a,** Schematic of in vitro transcription (IVT) method to generate base editor mRNA. Base editor constructs are cloned into an IVT production vector, amplified and poly-T tailed in a single PCR step, and then *in vitro* transcribed using modified capping and dNTPs to improve stability and reduce immunogenicity. **b,** Representative electropherogram trace of in vitro transcribed base editor mRNA indicating high purity and appropriate size of the mRNA products. **c,** Single- and multi-plex base editing of *B2M* and *CD2* with sgRNAs designed to generate gene knockout as described in Figure 1. **d,** Knockout editing of CD2 with lentiviral integration of a CD2 knockout guide followed by electroporation of Cas9 mRNA. sgRNAs were delivered in a CROP-seq vector with mTurquoise reporter, and mTurquoise expression shown in this histotogram indicates sgRNA expression of Cas9 perturbed cells. **e,** Representative histograms demonstrating similar base editing efficiency of *B2M* in both CD4 and CD8 T cells using lentiviral guide integration followed by base editor mRNA electroporation. **f,** Cytosine base editor-mediated knockout of T cell receptor (TCRab) surface expression by introduction of an early stop codon in the *TRBC* locus.

**Extended Data Figure 2. Library design and screen preparation. a,** Distribution of sgRNAs by category in the ClinVar ABE library. For pathogenic, likely pathogenic, and variant of unknown significance (VUS) categories, sgRNAs are predicted to either make the exact edit present in ClinVar, or an alternative missense mutation in the same amino acid. **b,** Distribution of sgRNAs by category in the 12-gene tiling ABE library. **c,** Schematic of sgRNA library cloning and high-titer library lentivirus production for base editor screens. **d,** Schematic for generation of library base-edited T cells. T cells were first isolated from healthy human donors, transduced with high-titer sgRNA library pools that were cloned into the CROP-seq mTurquoise lentiviral vector, followed by electroporation with ABE mRNA 72 hours later (see Fig. 1h). **e,** Transduction efficiency of ClinVar base editor library in two independent human donors T cells, which was performed at an optimized multiplicity of infection to achieve high frequency of cells receiving only one sgRNA. **f,** Representative histograms of IL2RA (CD25) sort in base edited T cells. For each donor, cells were sorted to select the top and bottom 15% of CD25 expression.

**Extended Data Figure 3. Metrics for rigor and reproducibility of large-scale base editing screens. a,** Density plots showing LFC values of different categories of guides from the ClinVar library at Day 35 post-electroporation of the long-term expansion screen arm. Dotted line represents the bottom 5% of combined empty window and silent mutation controls. Indicated are the percentages of guides in each category falling below this threshold. All guides generating variants in *CD3D, CD3E, CD3G,* or *CD3Z* were binned into the “CD3 complex” category. The second donor from the screen is shown (in companion to Fig. 3a). SgRNA distributions from all long-term proliferation timepoints are compared to a “Day 0” control T cell population which was library-transduced but not electroporated with base editor. **b,** Scatter plot showing LFC values of negative control sgRNAs (including both empty window and silent mutations) in both donors from the ClinVar Library at Day 28 post-electroporation in the long-term expansion screen arm with no significant enrichment or depletion of negative control guides is detected. **c,** Distribution of ranked relative abundance (RRA) scores for gene-wise dropout analysis in the CD25 sort arm of the ClinVar library, across both donors. The top 5 negatively selected genes (in top 15% of CD25 expression compared bottom 15% of CD25 expression) are listed. **d,** Distribution of ranked relative abundance (RRA) scores for gene-wise dropout analysis in the CFSE sort arm of the ClinVar library, across both donors. The top 5 negatively selected genes (in highly proliferative compared to least proliferative, for CFSE sort). **e,** Lollipop plot showing LFC of sgRNAs tiling IL2RG in the 12-gene tiling screen, mapped to the canonical IL2RG isoform. Selected sgRNAs generating clinically identified immunodeficiency variants, or alternative mutations at amino acids with known clinical variants, are annotated. Timepoint shown is Day 26 post-nucleofection in the long-term expansion screen arm. The second donor from the screen is shown (in companion to Fig. 3c). **f,** Scatter plot showing LFC of selected sgRNAs generating mutations in *LCK, SOS1,* and *PTPRC*. Timepoint shown is Day 28 post-electroporation in the ClinVar long-term expansion screen arm. Selected variants (e.g. LCK Y505C) were used for downstream functional validation experiments. **g,** Lollipop plot for sgRNAs targeting *RHOA* in one representative donor in the ClinVar screen, mapped to the canonical isoform. Timepoint shown is Day 35 post-electroporation in the long-term proliferation arm of the screen. **h,** Timecourse line graphs of LFC of sgRNAs targeting *RHOA* in one representative donor in the long-term expansion arm of the ClinVar screen. **i,** Volcano plot showing enriched and depleted guides in the CFSE lo vs hi proliferation sort. Positive LFC indicates enrichment of guides in the highly proliferative (CFSE lo) compared to least proliferative (CFSE hi) populations. One representative donor is shown (in companion to Fig. 3e). FDR cutoff <0.05. **j,** Volcano plot showing enriched and depleted guides in the CD25 hi vs lo proliferation sort. Positive LFC indicates enrichment of guides in the CD25hi compared to CD25lo populations. FDR cutoff <0.05. One representative donor is shown.

**Extended Data Figure 4. Screen results for genes undergoing tiled base editing. a-e,** Lollipop plots for genes targeted in the 12-gene tiling screen, with sgRNA editing positions mapped to the canonical isoform. Timepoint shown is Day 26 post-electroporation in the long-term expansion screen arm.

**Extended Data Figure 5. Screen results for additional genes undergoing tiled base editing. a-e,** Lollipop plots for genes targeted in the 12-gene tiling screen, with sgRNA editing positions mapped to the canonical isoform. Timepoint shown is Day 26 post-nucleofection in the long-term expansion screen arm.

**Extended Data Figure 6. Enrichment and structure-function relationship of variants promoting T cell proliferation. a,** Lollipop plot showing LFC of variants produced base editing screens and enriched following repeated stimulation. Indicated are selected variants and their amino acid positions across the gene product of PIK3CD (p110δ) and its highlighted domains. Representative timepoint shown is 15 days post-electroporation. **b,c,** Timecourse line graphs of LFC of sgRNAs targeting PIK3CD in both donors in the long-term expansion arm of the ClinVar screen. **d,e,** Lollipop plots for sgRNAs targeting AKT1 across both donors in the ClinVar screen, mapped to the canonical isoform. Timepoint shown is Day 35 post-electroporation in the long-term proliferation arm of the screen. **f,g,** Timecourse line graphs of LFC of sgRNAs targeting AKT1 in both donors in the long-term expansion arm of the ClinVar screen. **h,** Structure and position of mutations in AKT1. (right) Overall predicted structure of AKT1 (blue) and mutated residues (red). (left) Wild-type (WT) and mutated (mut) residues (red). D323G is predicted to localize next to L14 (dark blue). **i,j,** Lollipop plots for sgRNAs targeting LCK across both donors in the 12-gene tiling screen, mapped to the canonical isoform. Timepoint shown is Day 26 post-electroporation in the long-term proliferation arm of the screen. **k,** Structure and position of mutations in LCK. (top) Overall predicted structure of LCK (blue) and mutated residues (red). (bottom) Wild-type (WT) and mutated (mut) residues (red).

**Extended Data Figure 7. Design and results of co-culture experiments to improve T cell mediated lysis using variants identified in base editing screens. a,** Schematic for virus-free engineering and rapid expansion of NY-ESO-1 specific primary human T cells. **b,** Representative flow cytometry dot plot of NY-ESO-1 TCR-engineered T cells stained with TRAC antibody and NY-ESO-1 dextramer prior to flow sorting. **c,** Viable count relative to initial timepoint for A375-dsRed cells cultured at varying effector:target ratios of NY-ESO-1 specific T cells for 48 hours. aMHC = MHC class I blocking antibody, which blocks peptide-MHC-specific activation of T cells. n=3 replicates. **d,** Representative flow cytometry histograms of ABE- or CBE-mediated knockout of *B2M* or *CD2* in NY-ESO-1 specific T cells using sgRNAs shown in Figure 1. **e**, Mean fluorescence intensity (MFI, y axis) of TNFα expression in NY-ESO-1-TCR T cells with indicated base edits (x axis) after 8 hours of co-culture with A375-dsRED. One-way ANOVA **f**, Frequency (in %, y axis) of Granzyme B expression in NY-ESO-1-TCR T cells with indicated base edits (x axis) after 8 hours of co-culture with A375-dsRED. **g,** MFI (y axis) of Granzyme B expression in NY-ESO-1 specific T cells with indicated base edits (x axis) in the presence or absence of aMHC after 8 hours of co-culture with A375-dsRED. **h**, Flow cytometric evaluation of frequency (y axis) and intensity (x axis) of Granzyme B expression in NY-ESO-1 specific T cells with indicated genotypes from experiment shown in (g). **i,** Relative change in cell numbers of A375-dsRed 48 hours after co-culture with NY-ESO-1 variant T cells (effector-to-target ratio 0.5:1) from an independent experiment. n=3 replicates. **j,** Counts of viable A375-dsRed cells (y axis) over time (x axis, in hours) plated either alone or co-cultured at an effector-to-target ratio of 0.5:1 with NY-ESO-1 specific T cells harboring indicated mutations. T cells were added 4 hours after initial plating and imaging was performed using an Incucyte instrument at approximately 1-hour intervals. n=3 replicates. One-way ANOVA with Dunnett’s test for multiple comparisons (panels *c, e, f, g, i*).

## EXTENDED DATA TABLE LEGENDS

**Extended Data Table 1:** Genes selected for ClinVar screen, with STRING clustering by functional interactions.

**Extended Data Table 2:** List of essential genes selected for screen positive controls.

**Extended Data Table 3:** MAGECK Test outputs for all ClinVar screen comparisons.

**Extended Data Table 4:** MAGECK Test outputs for all 12-gene tiling screen comparisons.

**Extended Data Table 5:** All PCR oligos used in this study.

**Extended Data Table 6:** All sgRNA sequences used in this study.

**Extended Data Table 7:** All cloning oligos used in this study.

**Extended Data Table 8:** All sanger sequencing oligos used in this study.

