## Supplemental Figures for "Massively parallel base editing screens to map variant effects on anti-tumor hallmarks of primary human T cells"

### Extended Data Figure 1

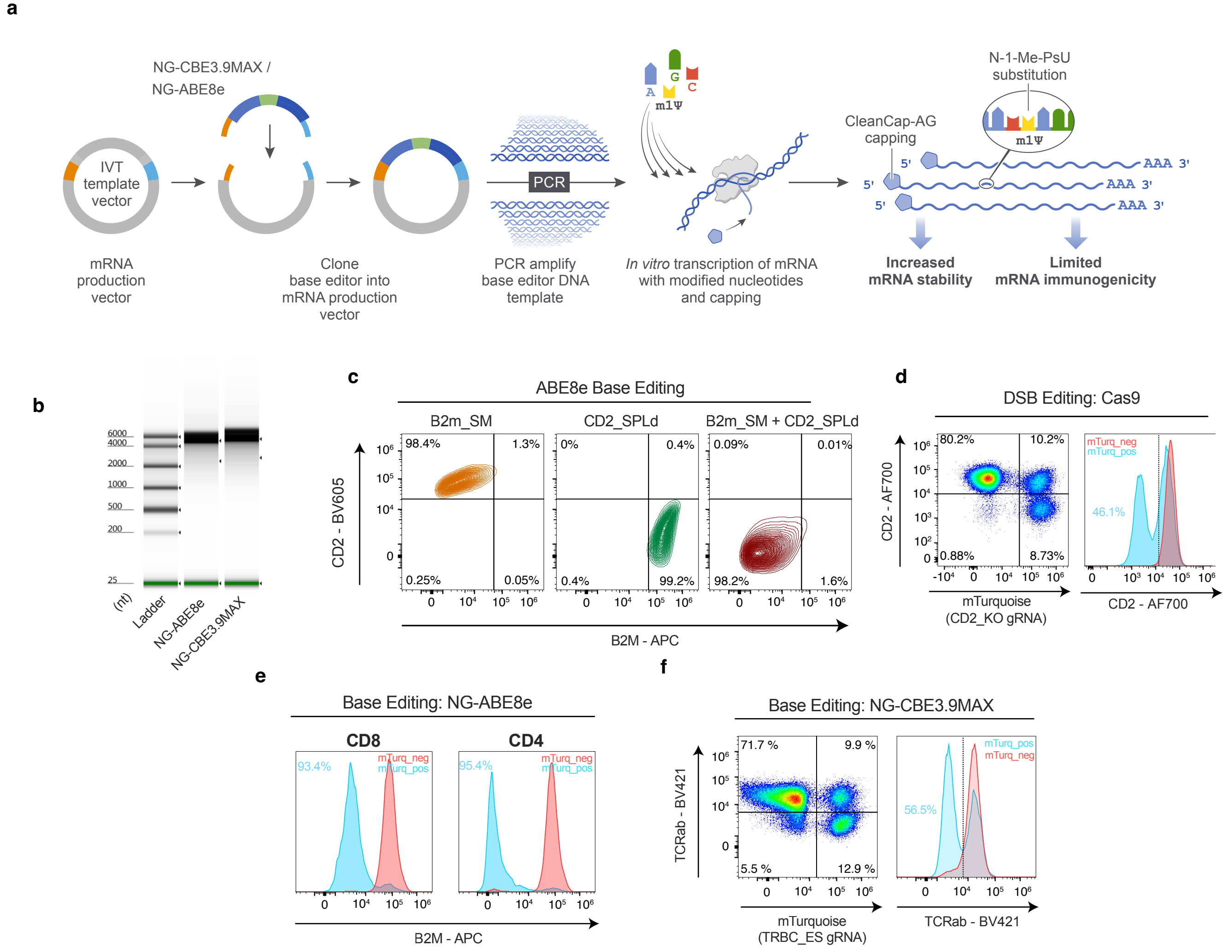

### Extended Data Figure 2

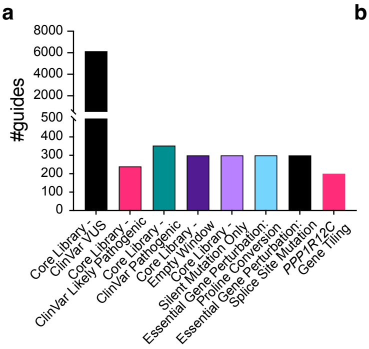


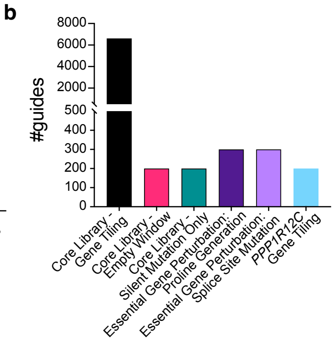


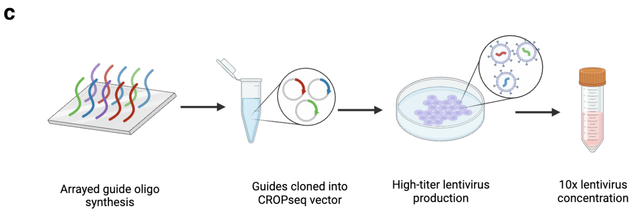


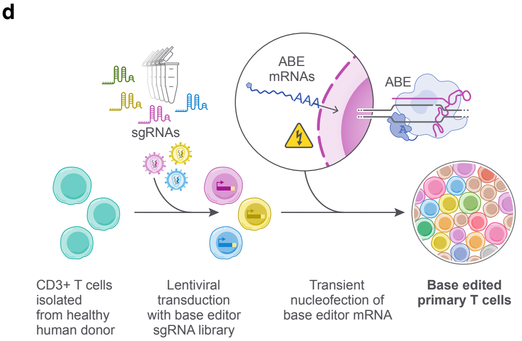


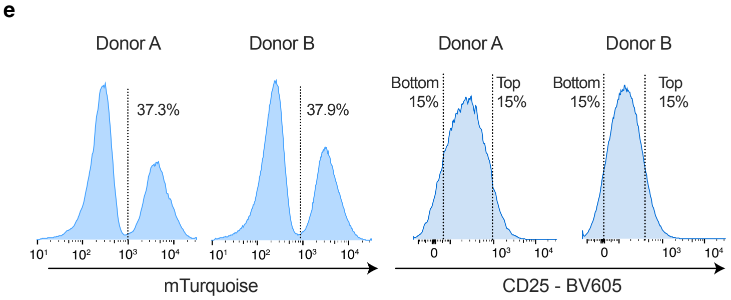

**Extended Data Figure 3**

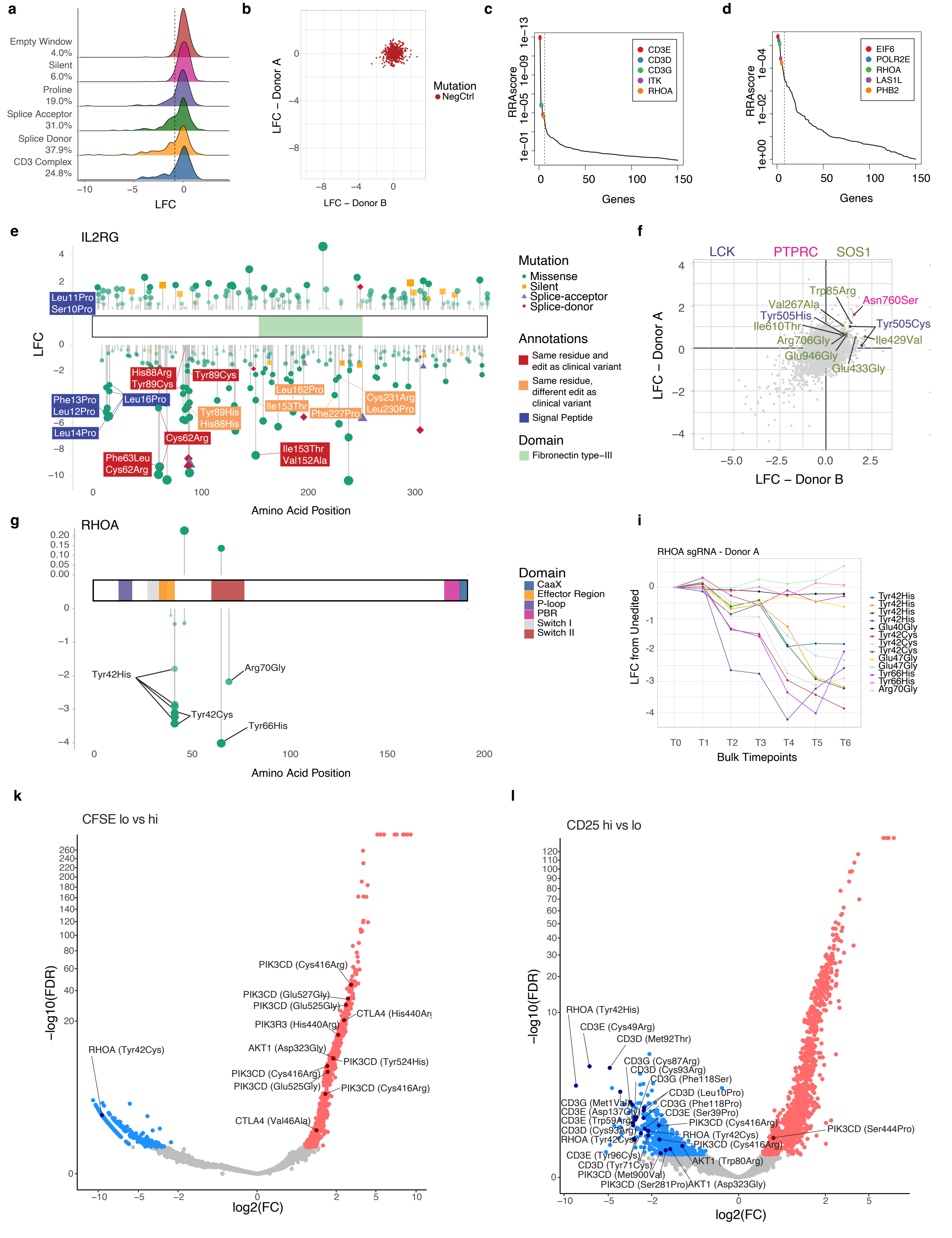

### Extended Data Figure 4

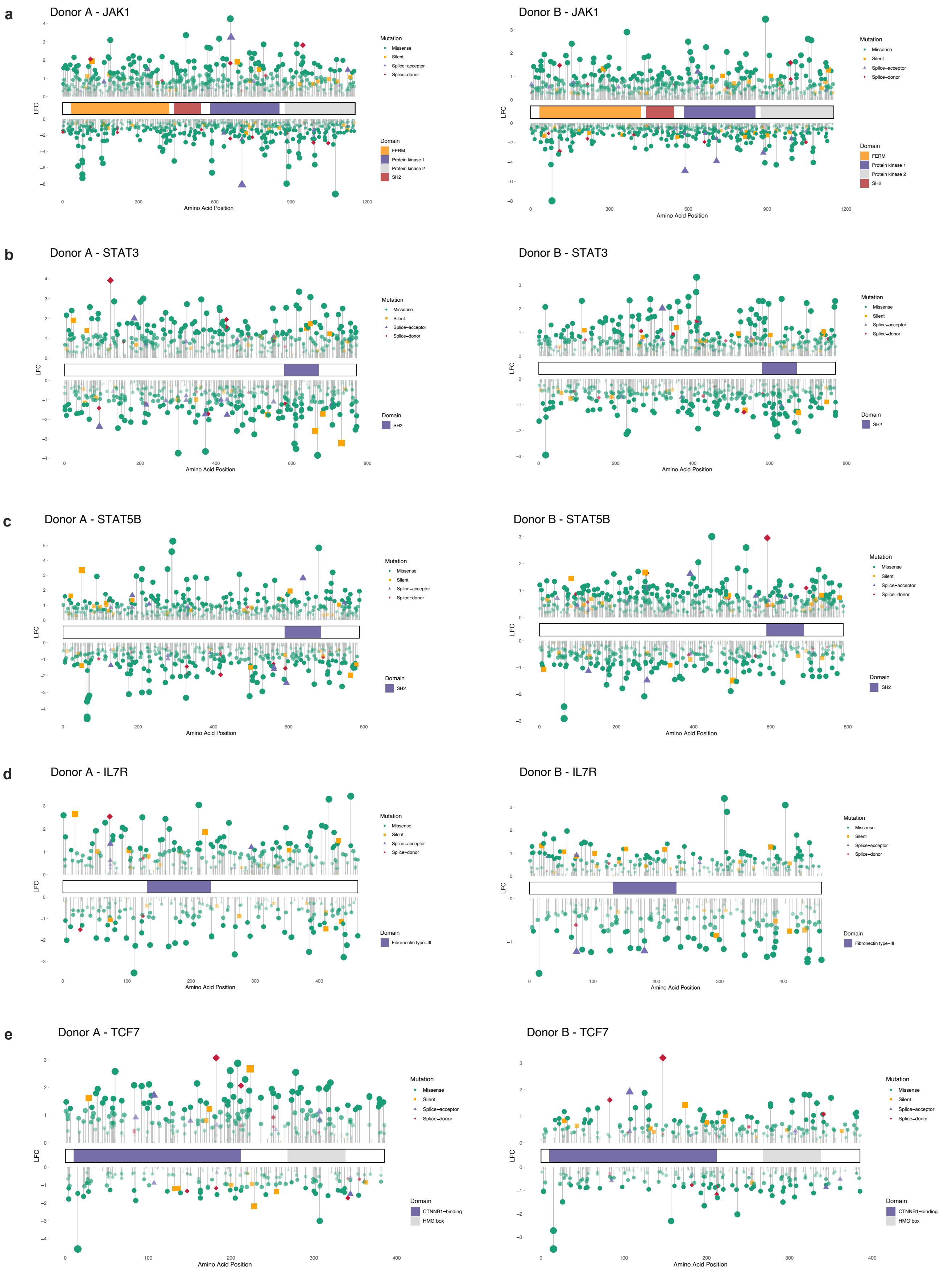

### Extended Data Figure 5

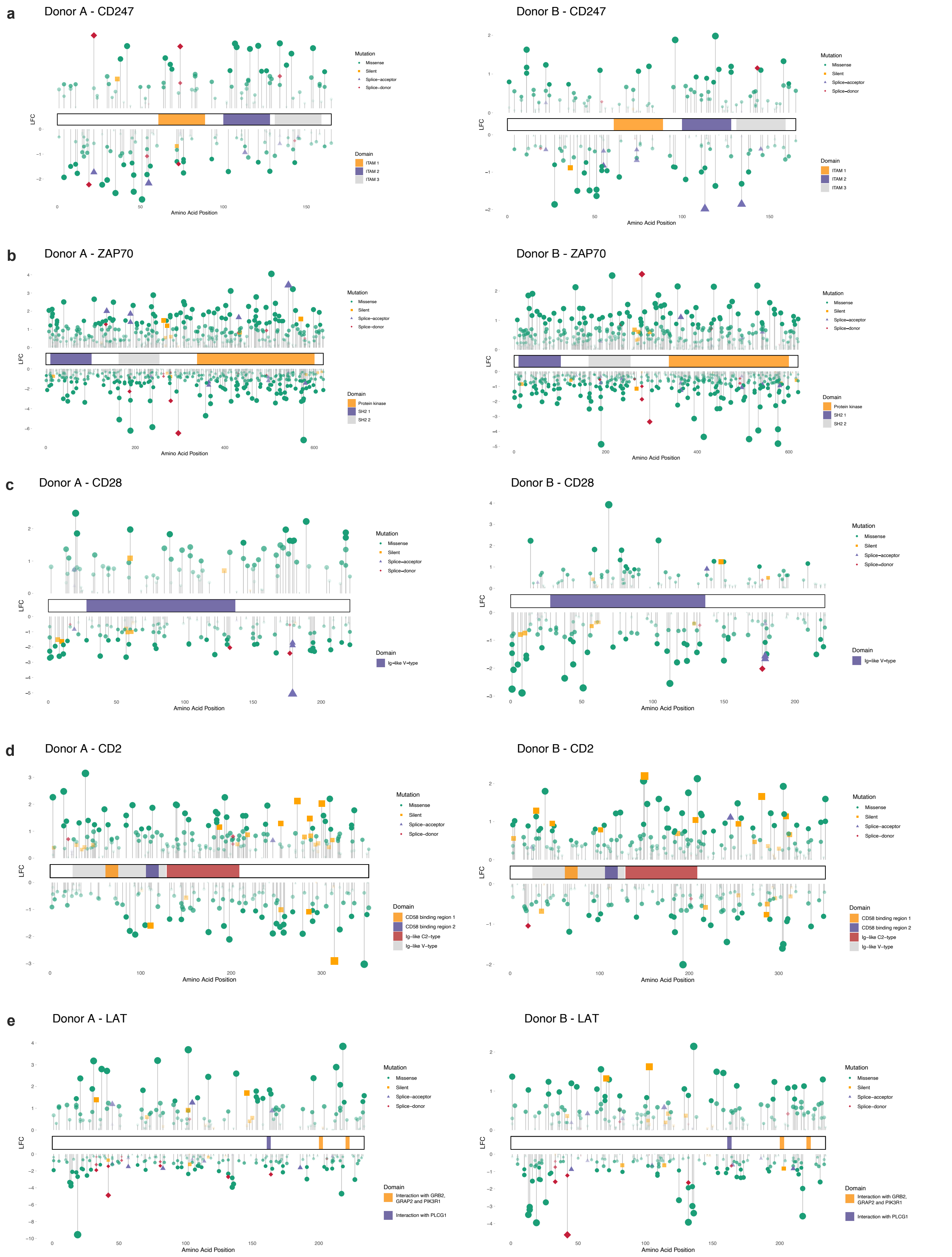

### Extended Data Figure 6

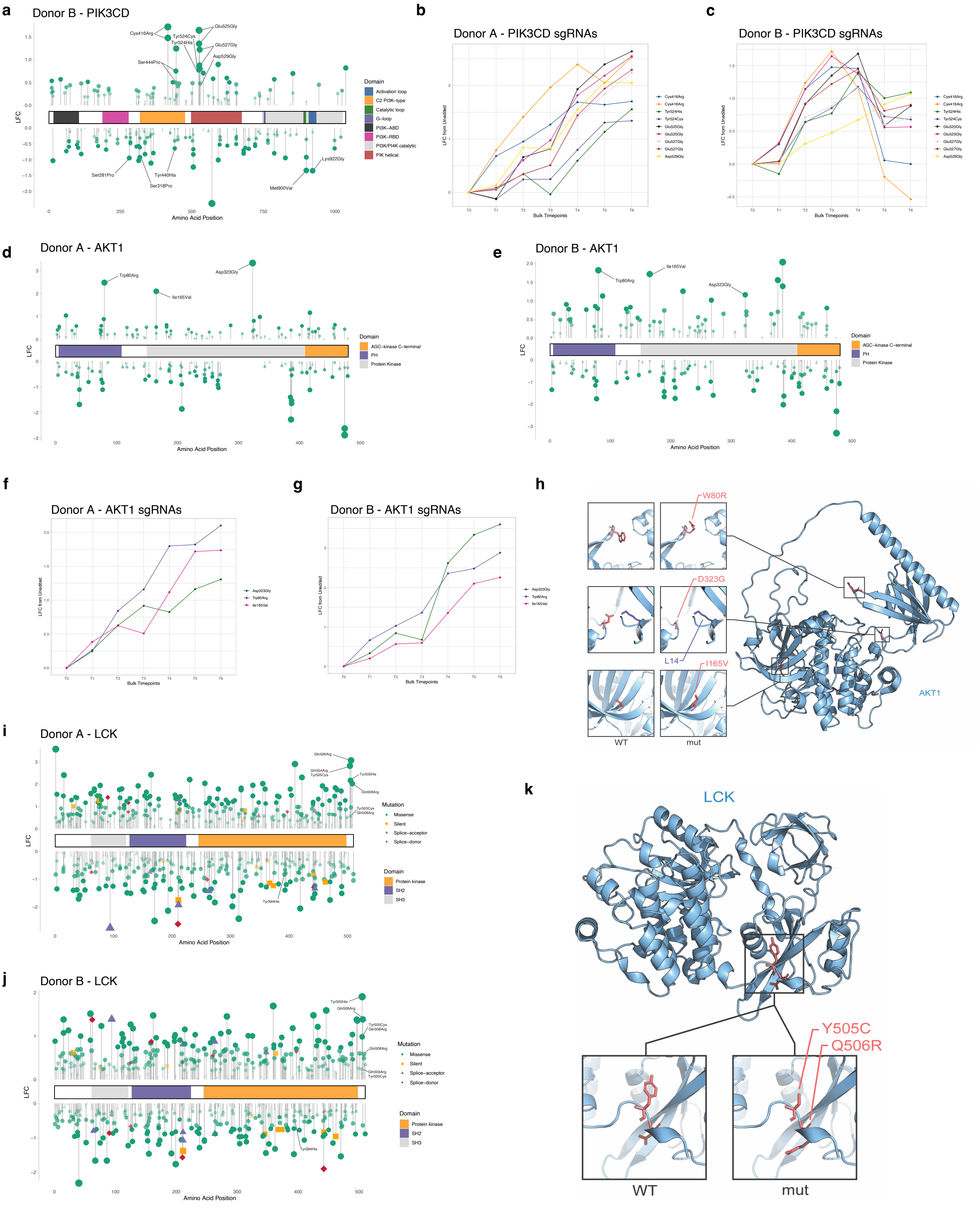

### Extended Data Figure 7

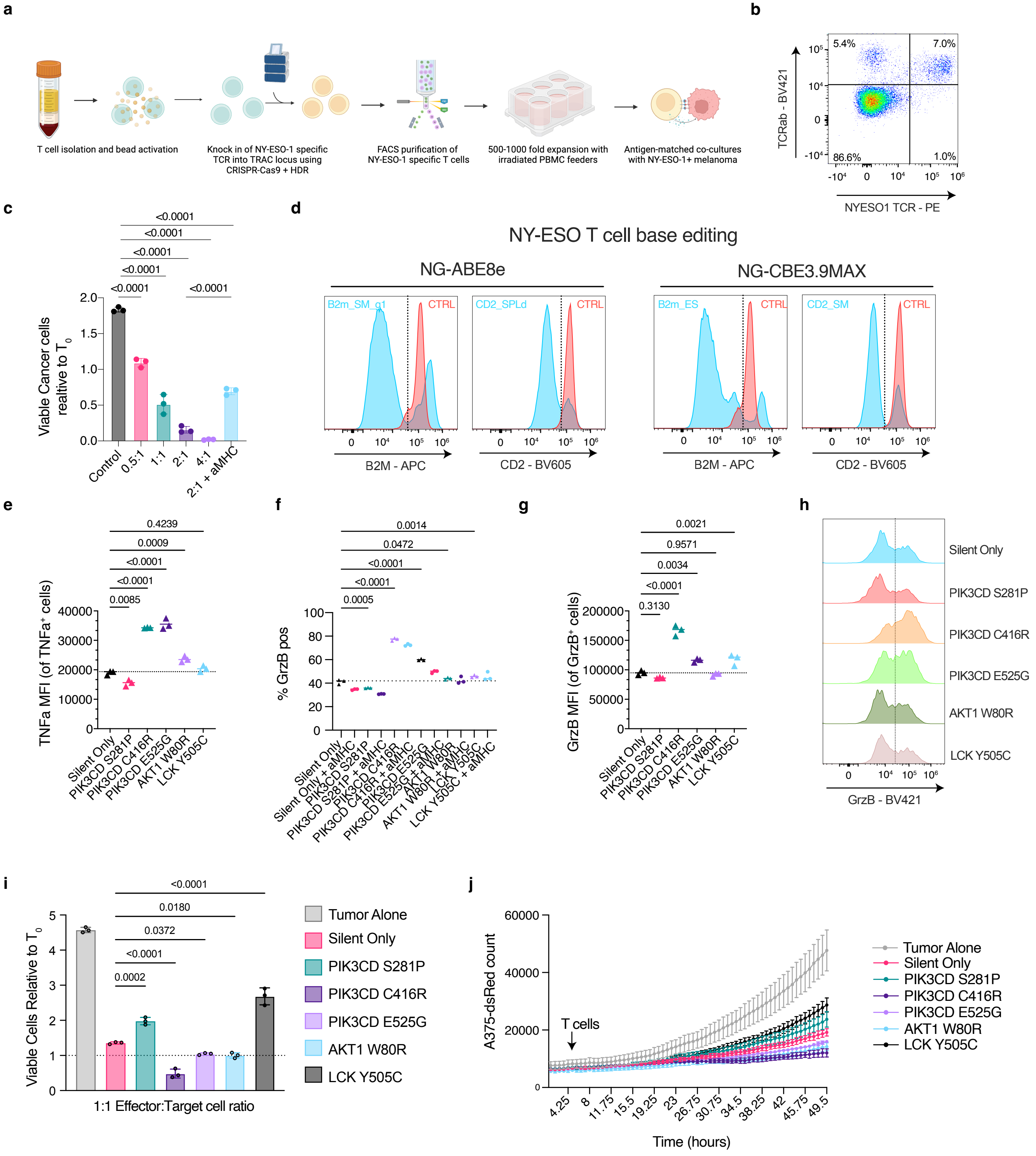
